# Modular subgraphs in large-scale connectomes underpin spontaneous co-fluctuation “events” in mouse and human brains

**DOI:** 10.1101/2023.05.17.538593

**Authors:** Elisabeth Ragone, Jacob Tanner, Youngheun Jo, Farnaz Zamani Esfahlani, Joshua Faskowitz, Maria Pope, Ludovico Coletta, Alessandro Gozzi, Richard Betzel

**Affiliations:** Neuroscience Program, Oberlin College, Oberlin, OH 44074; Cognitive Science Program and School of Informatics, Computing, and Engineering, Indiana University, Bloomington IN 47401; Department of Psychological and Brain Sciences and Cognitive Science Program, Indiana University, Bloomington IN 47401; Stephenson School of Biomedical Engineering, The University of Oklahoma, Norman, OK 73019; Section on Functional Imaging Methods, National Institute of Mental Health, Bethesda, MD 20892; Program in Neuroscience and School of Informatics, Computing, and Engineering, Indiana University, Bloomington IN 47401; Fondazione Bruno Kessler, Trento, Italy; Functional Neuroimaging Lab, Istituto Italiano di Tecnologia, Center for Neuroscience and Cognitive Systems, Rovereto, Italy; Department of Psychological and Brain Sciences, Cognitive Science Program, Program in Neuroscience, and Network Science Institute, Indiana University, Bloomington IN 47401

## Abstract

Previous studies have adopted an edge-centric framework to study fine-scale dynamics in human fMRI. To date, however, no studies have applied this same framework to data collected from model organisms. Here, we analyze structural and functional imaging data from lightly anesthetized mice through an edge-centric lens. We find evidence of “bursty” dynamics and events – brief periods of high-amplitude network connectivity. Further, we show that on a per-frame basis events best explain static FC and can be divided into a series of hierarchically-related clusters. The co-fluctuation patterns associated with each centroid link distinct anatomical areas and largely adhere to the boundaries of algorithmically detected functional brain systems. We then investigate the anatomical connectivity undergirding high-amplitude co-fluctuation patterns. We find that events induce modular bipartitions of the anatomical network of inter-areal axonal projections. Finally, we replicate these same findings in a human imaging dataset. In summary, this report recapitulates in a model organism many of the same phenomena observed in previously edge-centric analyses of human imaging data. However, unlike human subjects, the murine nervous system is amenable to invasive experimental perturbations. Thus, this study sets the stage for future investigation into the causal origins of fine-scale brain dynamics and high-amplitude co-fluctuations. Moreover, the cross-species consistency of the reported findings enhances the likelihood of future translation.

## INTRODUCTION

A growing body of literature has shown that co-ordinated brain activity supports ongoing neural, behavioral, and cognitive processes. These activity patterns are constrained by the brain’s underlying structural connectivity (SC), whose network configuration organizes brain activity into cohesive and correlated patterns–i.e. functional connectivity (FC) [1, 2].

The correlation structure of neural activity is not static; rather it fluctuates from moment to moment [3, 4]. There exist many techniques for tracking these rapid fluctuations, including sliding window [5] and kernel-based methods [6]. However, these approaches generate estimates of time-varying FC (tvFC) that are temporally imprecise. That is, the estimate of FC at time *t* depends not only on the state of the brain at that instant, but also on its state at nearby time points [7].

Recently, we built upon existing frameworks for estimating changes in functional network structure [8–12] to develop a technique–referred to as “edge time series– for tracking instantaneous co-fluctuations between pairs of brain regions–i.e. network edges [13, 14]. Using this approach, we found evidence of global “events”–brief periods of high-amplitude and brain-wide co-fluctuation. We showed that events express subject-specific information [15, 16], can be used to approximate static FC and enhance brain-behavior associations [13, 17], and can be partitioned into a clusters of repeating patterns [15, 18], whose relative frequency may be linked to variation in endogenous hormone fluctuations [19].

Developing insight into the origins of events is the subject of ongoing work [20–22]. First, several studies have shown that event timing is correlated across individuals during movie-watching [13, 23, 24], suggesting that naturalistic audiovisual stimuli can initiate cascades of neuropsychological processes that act to support the appearance of events in fMRI time series. Second, other studies have demonstrated that events can occur spontaneously in networked dynamical systems [25]. For instance, Pope *et al*. [26] found that when the underlying constraint matrix–i.e. structural connectivity–exhibits modular structure, events naturally occur and that their topography aligns closely with the boundaries of anatomical modules. In aggregate, these findings suggest that event-like activity has both a cognitive underpinning, but can also emerge due to modularity in the underlying system structure.

Despite the fact that many studies have applied this “edge-centric framework” to human imaging data, to our knowledge it has never been extended to non-human data. Such an extension could prove particularly useful, as the rich set of (sometimes invasive) perturbations [27, 28] that can be applied to non-human subjects could help address open controversies and questions surrounding the origins of events and the importance of time-varying coupling between brain areas. Additionally, few empirical analyses have examined the link between structural connectivity and high-amplitude edge-centric co-fluctuations (though see [25]).

Here, we take the initial step in that direction, applying event-detection to two datasets: first to fMRI BOLD data acquired from 18 anesthetized mice and subsequently to a large human cohort (Human Connectome Project [29]). In line with previous studies, we find evidence of events and show that events are highly predictive of static FC and can be grouped into hierarchically related co-fluctuation patterns. As demonstrated in Sporns *et al*. [30], each cluster centroid results in a bipartition of the brain into two disjoint sets of nodes, one positively co-fluctuating and the other negatively co-fluctuating. Finally, we show that the bipartitions are underpinned by highly modular sub-networks in SC, positing an anatomical basis for opposed co-activity. Further, we replicate all of these findings using human functional imaging data. Collectively, our findings set the stage for future, more targeted and hypothesis-driven investigations into the anatomical underpinnings of fine-scale, network-level co-fluctuations.

## RESULTS

In this paper we sought to replicate several key findings initially made using data acquired from human subjects in mouse imaging data. First, we wanted to show that on a per-frame basis, high-amplitude events better recapitulate static FC than middle-/low-amplitude frames. Second, we wanted to show that events could be meaningfully partitioned into a small set of recurring “states” or “event clusters”. Lastly, we wanted to explore, using empirical data, the link between anatomical connectivity and high-amplitude events. As part of this exploration we include a second (human) imaging dataset, allowing us to replicate findings initially made using mouse data with a much larger data in which anatomical connectivity was mapped non-invasively, thereby opening up the possibility of future translational studies. In this section, we report the results of these analyses.

### High-amplitude co-fluctuations recapitulate static FC

One of the first observations made using edge time series was that, with only a small subset of high-amplitude frames–putative “events”–it was possible to accurately reconstruct static FC [13]. These previous observations were made using data recorded from awake human participants. Here, we assess whether a similar effect is evident when we apply edge time series to functional imaging data acquired from anesthetized mice.

Our procedure for testing this hypothesis included a series of post-processing analysis steps. First, we transformed fMRI BOLD time series from *N* = 182 parcels (Fig. 1a) into edge time series. This procedure involved standardizing (z-scoring) each time series and, for each of the *N* (*N* − 1)/2 = 16471 pairs, calculating their framewise product (Fig. 1b). The result is a co-fluctuation or “edge time series” for every pair of nodes whose elements encode the timing, amplitude, and sign of interregional co-fluctuations. Notably, the temporal average of a given edge time series is exactly the bivariate product-moment correlation–i.e. FC (Fig. 1c).

**Figure 1.**
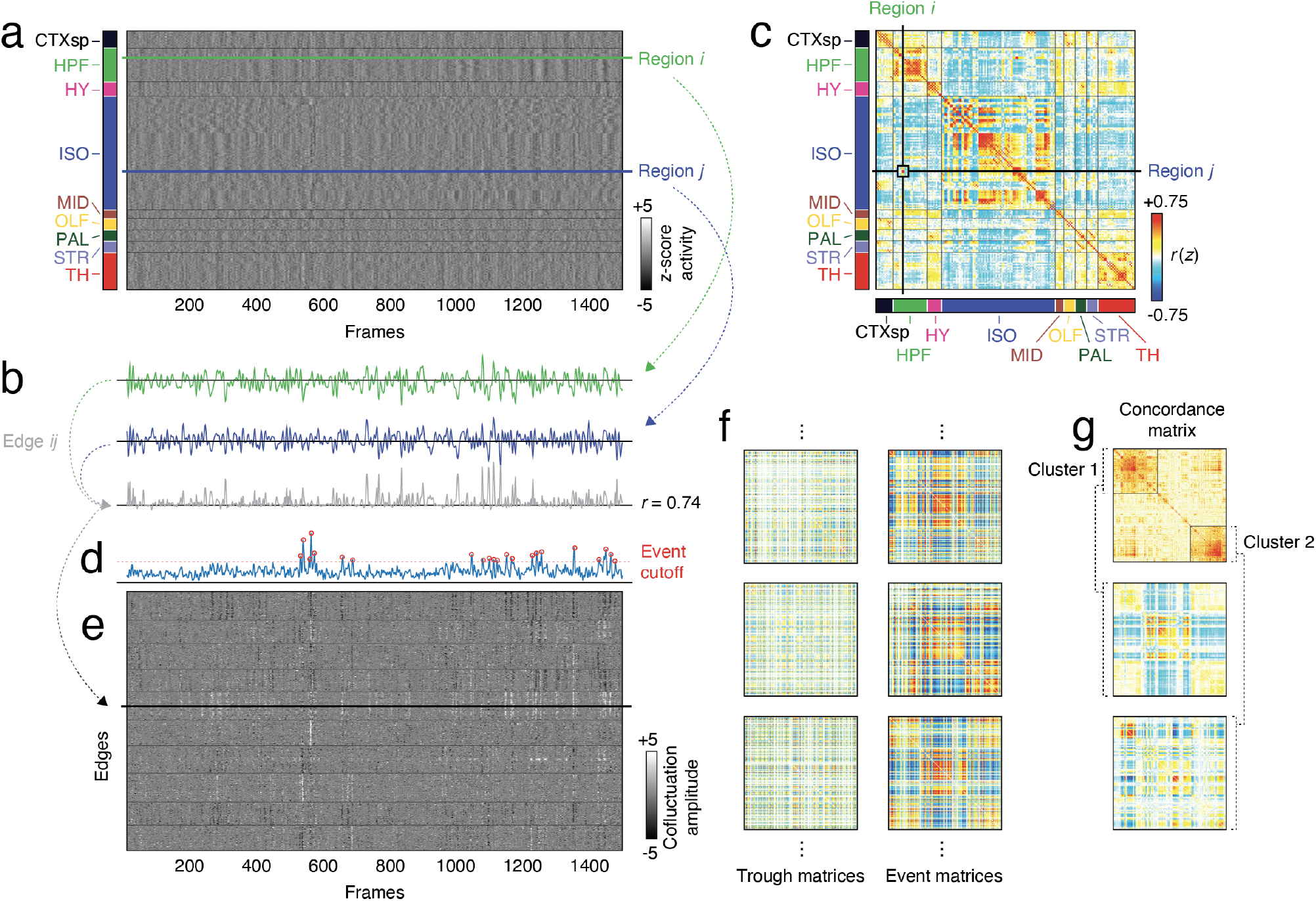
Schematic illustrating edge time series construction and clustering. (*a*) Parcel time series. (*b*) Functional connectivity is usually estimated as the correlation between pairs of parcel times. Correlation can be thought of as the normalized sum of the element-wise product between pairs of parcel time series. (*c*) That value is entered into the symmetric functional connectivity/correlation matrix. Omitting the normalized summation step yields a “co-fluctuation time series” whose elements correspond to the framewise product between standardized parcel time series. This procedure can be repeated for all pairs of time series, generating a matrix of co-fluctuation or “edge time series” (see panel *e*). (*d*) The global amplitude of co-fluctuations can be calculated as the root mean square across all edge time series at every frame. (*f*) The set of all edge time series at a given frame can be reshaped into the upper triangle elements of a “co-fluctuation matrix”. The highest amplitude frames that exceed a statistical threshold can be labeled “events” and analyzed further. (*g*) Event co-fluctuation patterns can be aggregated across animals and clustered into putative states.

To detect events, we analyzed all edge time series collectively (Fig. 1e). At each frame we calculated the global co-fluctuation amplitude as the root mean square (RMS) across all edge time series. This step yielded a single time series whose peaks could be detected easily (using MATLAB’s `findpeaks.m` function). The amplitudes of these peaks were compared against a null distribution generated by independently circularly shifting parcel time series, recalculating edge and RMS time series, and aggregating peaks of the null RMS time series across 100 runs. Empirical peaks whose amplitude was significantly greater than that of the null distribution were categorized as events (statistical significance was established at the subject level; critical *p*-value adjusted to maintain a false-discovery rate fixed at *q* = 0.05).

Following event detection and separately for each sub-ject, we averaged the co-fluctuation patterns expressed during each event and calculated the similarity (correlation) of this pattern with static FC. Across all animals, the number of events was fewer than the total number of non-event peaks (peaks in the RMS time series that did not reach the statistical criterion for being considered an event), troughs (local minima in the RMS time series), and the total number of frames (by definition). To assess whether events were more similar to these other categories and to control for differences in the number of frames associated with each category, we randomly subsampled a number of frames equal to the number of events from within each category, averaged the co-fluctuation patterns across these frames, and calculated its similarity with respect to static FC (we performed 100 repetitions of this subsampling procedure; see Fig. 2a for an example of a single set of subsamples and Fig. 2b for the results from 100 sub-samples from one subject).

**Figure 2.**
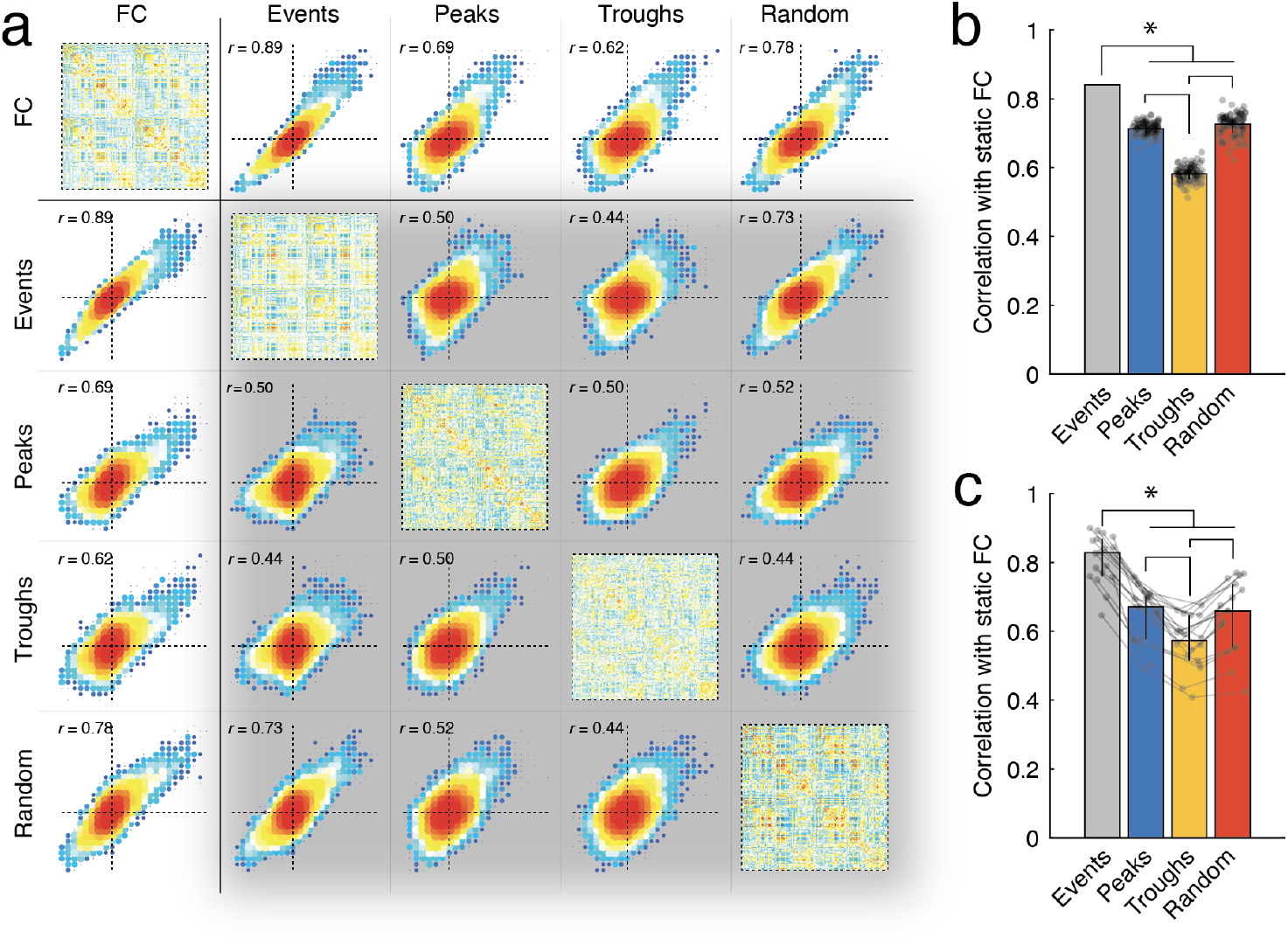
Differential correlation between frame categories and static FC. Previous studies have documented a strong correspondence between static FC and high-amplitude frames (events). Here, we compare four categories of frames: high-amplitude peaks (*Events*), peaks that are not considered high-amplitude (*Peaks*), low-amplitude frames (*Troughs*), and random selections of frames (*Random*). (*a*) The diagonal shows example co-fluctuation matrices from each of the four categories as well as static FC. The off-diagonal blocks show example scatterplots between each pair of categories. Matrices and scatterplots depicted here come from a single mouse and for the sub-sampled categories, a single sub-sample. (*b*) Example correlations from a single subject over 100 random sub-samples from within each category. Each sub-sample contained the same number of frames as the number of detected events. (*c*) Median correlations aggregated across all 18 mice. Lines connect data points from the same mouse. Vertical lines represent *±*1 standard deviation.

We then averaged similarity scores across the 100 repetitions for each subject and compared the mean similarity across the four categories of frames: events, non-event peaks, troughs, and random sets of frames (Fig. 2c). We observed that across all subjects, events were significantly more similar to static FC than other frame categories on a per frame basis. The mean similarity of non-event peaks was not significantly different from random samples, while all other frames were significantly more similar to FC than troughs were to FC (false discovery rate fixed at *q* = 0.05 and critical *p*-value adjusted accordingly).

Note that while these observations are consistent with previous findings [13, 15], other studies have reported that the highest-amplitude frames may not be optimal in terms of recapitulating static FC (see for example [16, 22]). Rather, those studies find that the second highest amplitude bin outperforms the highest. Why might this be? We investigate this question in Fig. S1. We compared the “binning of all frames” approach of Cutts *et al*. [16] and Ladwig *et al*. [22] with “peak binning”, an approach that is more similar to what we report in Fig. 2. We find that using the “all frames” approach we could replicate the effect described by Cutts *et al*. [16] and Ladwig *et al*. [22]. With this approach, the extreme bins – the highest- and lowest-amplitude frames – are comprised of co-fluctuation patterns that are temporally proximal to one another. This is likely due to the strong serial correlation of the fMRI BOLD signal, the relative infrequency of events, and the increased number of frames necessary to rise to a high-amplitude event relative to lower-amplitude peaks [18]. In contrast, the “peak binning” procedure exhibited no such bias. Therefore, a possible explanation for the apparent superior performance of the second highest amplitude bin is that it tends to sample the entire scan session better than the highest-amplitude bin.

### High-amplitude events can be sub-divided into recurring network states

Previous studies have shown that high-amplitude and network-level events can be clustered into putative states on the basis of their topographic similarity to one another [15, 18, 19]. It is unclear whether the same is true for the mouse edge time series data analyzed here.

To address this question, we followed the analysis pipeline from Betzel *et al*. [18]. Briefly, this involved aggregating event co-fluctuation patterns across all subjects, calculating the similarity (Lin’s concordance) between all pairs of events (Fig. 3a), and recursively applying modularity maximization (coupled with consensus clustering and a statistical criterion for terminating the recursion) to obtain a hierarchy of statistically significant event clusters (Fig. 3b,c).

**Figure 3.**
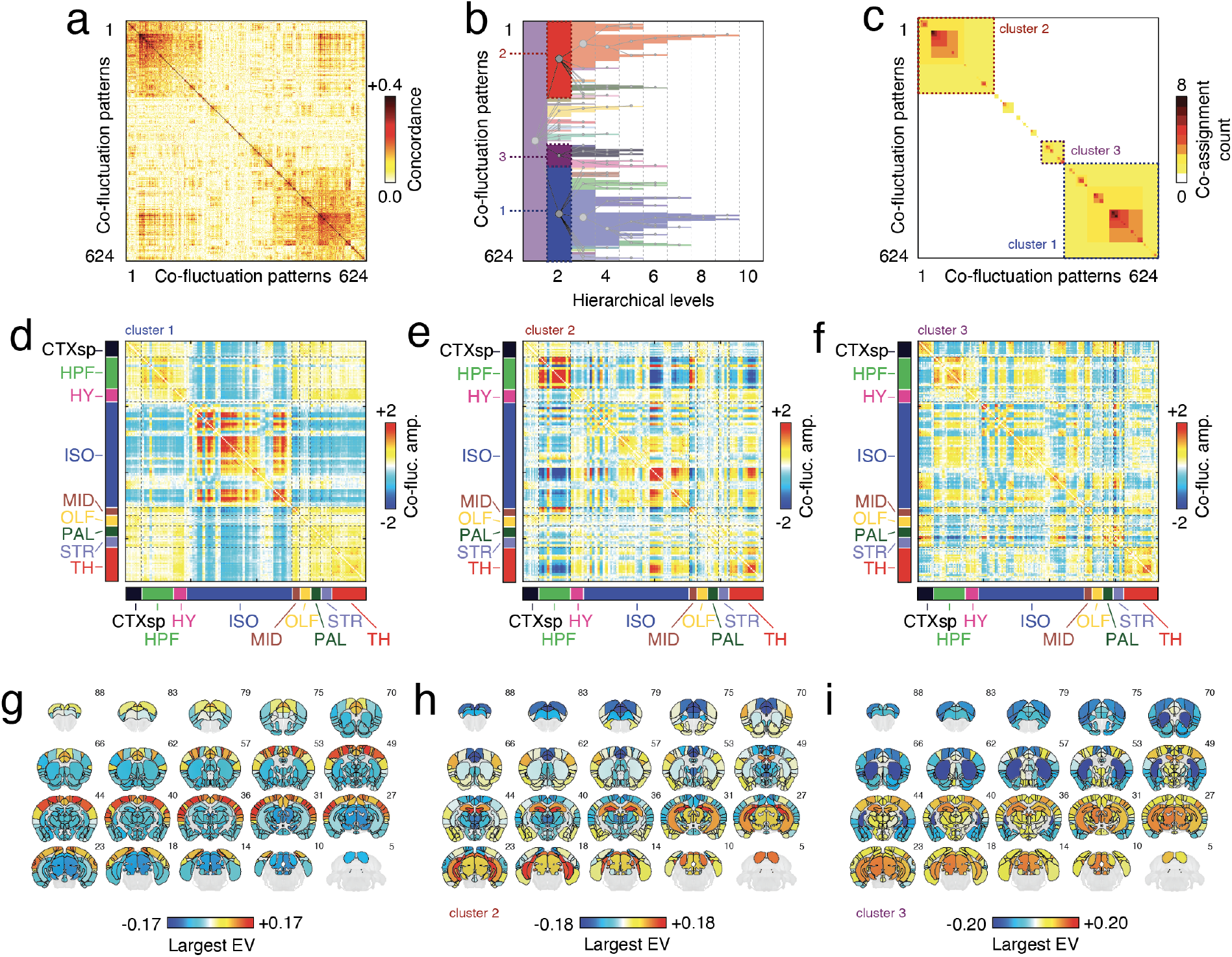
Hierarchical clustering of high-amplitude co-fluctuations. (*a*) Concordance matrix. Rows and columns represent events co-fluctuation matrices aggregated across subjects. (*b*) Hierarchical clustering of co-fluctuations patterns. Gray circles represent clusters and lines indicate parent-child relationships. (*c*) Co-assignment matrix (counts) from hierarchical clustering. Panels *d, e*, and *f* represent centroids for the three large clusters detected at hierarchical level 2. Panels *g, h*, and *i* depict largest eigenvector of each matrix projected back into anatomical space.

Across subjects, we identified 624 putative “events” (34.7 ± 10.2 events per animal out of 1500 frames; maximum of 49; minimum of 16). The hierarchical clustering procedure grouped the corresponding co-fluctuation patterns into ten hierarchical levels, eight of which were non-trivial (the first and last levels corresponded to a single community and no communities, respectively). For brevity, we focus on the second hierarchical level (the first non-trivial level), which exhibited a total of 12 distinct event clusters, three of which stood out as they collectively accounted for 39.3%, 31.7%, and 9.5% of all co-fluctuation patterns (the next largest cluster contained 5.1% of patterns).

The first event cluster was typified by opposed activation of isocortex with midbrain and the hippocampal formation (Fig. 3d,g; see Fig. S2 for details of anatomical labels). This pattern of connectivity has been described at length in mouse imaging literature as a murine analog of the default mode network [31, 32]. It also mirrors findings made in the human literature, where the largest event cluster also implicates default mode co-fluctuations [13, 15, 18, 19]. Cluster two was typified by strong co-fluctuations of the hippocampal formation, an areas thought to support memory formation and recall, with components of isocortex and thalamus, which have been implicated in attention and perception (Fig. 3e,h). Finally, cluster three involved strong opposed activity of regions in striatum, isocortex, and the cortical subplate with regions in the hippocampal formation, other isocortical parcels, and midbrain (Fig. 3f,i). Whereas clusters one and two are mostly refined across hierarchical levels, cluster three neatly splits into two sub-clusters in the third hierarchical level, the first of which emphasized opposed activity of the hippocampal formation with the cortical subplate, striatum, and pallidum, while the second emphasized opposed activity of thalamus with isocortex and to a lesser extent, the cortical subplate (see Fig. S3).

An alternative and complementary view of high-amplitude events can be obtained by considering their alignment with respect to functional systems obtained using data-driven techniques–e.g. clusters or “modules” derived from static FC. To this end, we performed a hierarchical decomposition of static FC, revealing multilevel network organization (Fig. S4). Interestingly, the co-fluctuation patterns associated with the event clusters described above neatly align with these static modules. For instance, the first event cluster corresponds to strong co-fluctuations of module 1 (M1) with the other three modules. The second and third event clusters correspond to opposed co-fluctuations of module 4 (M4) with modules 1 and 2 (M1; M2) and module 2 with, largely, the rest of the brain (Fig. S5). Note that we also recapitulate these findings using high-resolution near-voxel-level data (Fig. S6).

### High-amplitude co-fluctuation patterns reflect modular sub-divisions of mouse anatomical connectivity

In Pope *et al*. [26], the authors detected events in synthetic fMRI BOLD data generated by an anatomically constrained oscillator model. The co-fluctuation patterns associated with these events could be mapped back to the underlying anatomical network, specifically its modular structure. However, the relevance of anatomical modules for high-amplitude edge-level events has never been validated empirically (with human data or otherwise). Moreover, human structural connectivity derived from water-diffusion statistics–as used by Pope *et al*. [26]–exhibits some biases [35–37], including difficulty in accurately tracking interhemispheric fibers [38, 39], thereby making a direct empirical replication of Pope *et al*. [26] less likely.

Instead, we examine the high-amplitude co-fluctuations in mice and their structural underpinnings. Here, the murine connectome was invasively mapped using viral tracers and tract tracing techniques [40], thereby circumventing some of the limitations associated with tractography and diffusion MRI data. In addition, the invasive tracing technique allows for the mapping of directed connections, a feature not resolvable using water-diffusion methods.

Our strategy for comparing event co-fluctuations and anatomical connectivity deviated from that of Pope *et al*. [26], which depended upon a specific definition of anatomical modules. Instead, our approach was to recover the bipartition of network nodes into positively and negatively fluctuating groups associated with each event cluster centroid [30]. If we were to examine a single co-fluctuation pattern, its bipartition is defined unambiguously. However, for event cluster centroids, which reflect the mean over many co-fluctuation patterns, recovering the bipartition is not as straightforward and requires additional analysis. One possible solution is to apply clustering algorithms–e.g. modularity maximization–to the centroid networks (see **Materials and Methods**). Given an estimate of the bipartition, we then imposed this partition onto the network of structural connections and calculated the modularity that it induced [41]. We compared the observed modularity against two null models; one in which we randomly assign nodes to either group, destroying spatial autocorrelations, and another in which we approximately preserve the variogram–i.e. the spatial dependencies–of the original data [42].

Interestingly, we found that for the two largest event clusters, their induced modularity exceeded what was expected under both null models (1000 permutation tests; *p <* 10^*−*3^ for the independent permtuation model; maximum *p*-value of *p* = 0.03 for the geometry-preserving model; Fig. 4). For cluster three, the modularity was significantly greater than that of the independent permutation model (*p <* 10^*−*3^) but not significantly greater than the geometry-preserving model (*p* = 0.194). For event cluster one, these results were not dependent on the resolution parameter used define the cluster. However, for clusters two and three, there were select ranges where the geometry-preserving model achieved modularity scores consistent with what was observed in the empirical network, underscoring the role of geometry in constraining both the configuration of anatomical connections as well as functional patterns of activity/connectivity (Fig. S7). Interestingly, event co-fluctuation patterns induced greater structural modularity than non-event peaks (*p <* 10^*−*3^; Fig. S8), suggesting that the link between structural modularity and co-fluctuation patterns is uniquely strong for events. Indeed, this observation speaks to a more general relationship between co-fluctuation patterns and structural connectivity in which the correspondence between the two is greatest during periods of high-amplitude co-fluctuations (Fig. S9). As before, this effect holds with networks defined at finer spatial scales (Fig. S6).

**Figure 4.**
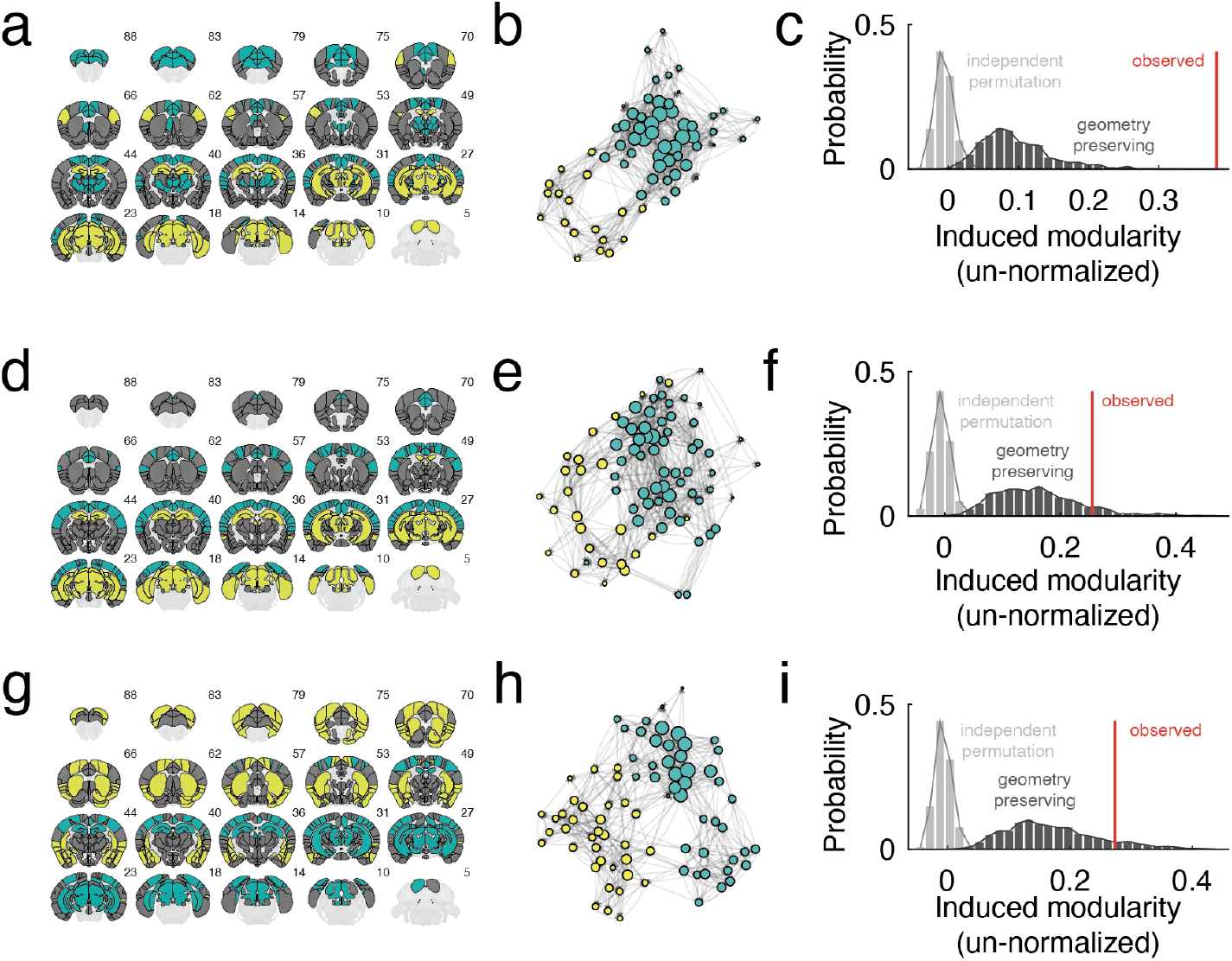
Linking high-amplitude events to structural connectivity. Panels *a, d*, and *g* represent bipartition communities for each of the three largest event cluster centroids in hierarchical level 2. Panels *b, e*, and *h* force-directed layouts of the induced sub-graph containing only nodes in either of the bipartition communities. Panels *c, f*, and *i* show the induced modularity of each sub-graph.

### Replicating event-module relationships using human imaging data

In the previous section, we found that whole-brain co-fluctuation patterns were underpinned by modular sub-divisions of the mouse connectome. It unclear whether a similar effect holds using human imaging data. Unlike the mouse connectome, the human connectome is typically reconstructed non-invasively from tractography and diffusion MRI. Although human connectome data has known limitations [35–37], its promise for translation and for understanding uniquely human neuropsychiatric disorders is greater. In this section, we replicate the main results from the previous two sections.

Briefly, our replication involved detecting events in resting-state data from the Human Connectome Project. We focused on a subset of the 100 unrelated participants that passed quality checks and motion exclusion criteria (see [14, 16]). For each subject and scan (95 subjects × 4 scans each), we performed event detection, aggregating event co-fluctuation patterns across individuals. This procedure resulted in 12854 events, which were subsequently partitioned hierarchically (Fig. 5a). As with the mouse data, we focused on the second hierarchical level, which yielded three large clusters (Fig. 5b,c,e,f,h,i). These cluster centroids were in line with those reported in other studies of event clusters [15, 18, 19]. Next, using a group-representative structural connectivity matrix [43], we calculated the modularity induced by each cluster. As in the previous sections, we compared the observed modularity against a null distribution generated under a permutation-based model in which nodes were randomly assigned to one of the two bipartition communities and a geometry-preserving “spin test” [33, 34]. In all cases, the observed modularity exceeded that of both null model (10000s permutations; *p <* 10^*−*4^). Note that in the supplement we also replicate using human imaging data the earlier finding that time-varying structure-function coupling depends on RMS (Fig. S10).

**Figure 5.**
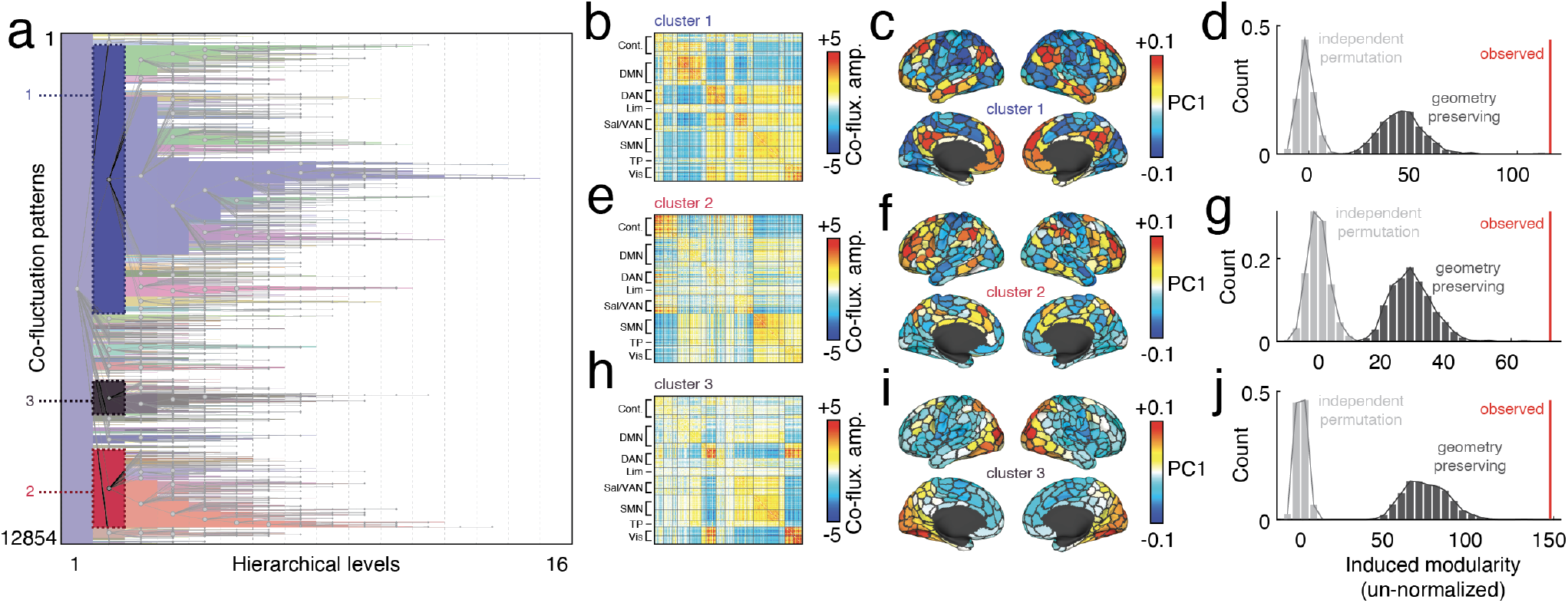
Replication of mouse findings using human MRI data. (*a*) Hierarchical clustering of 12854 event co-fluctuation patterns from 95 participants in the Human Connectome Project. Panels *b, c*, and *d* show cluster centroids for the three largest event clusters in hierarchical level 2, their projection onto the cortical surface, and the structural modularity induced by a bipartition derived from the co-fluctuation matrix. Here, we include a spatially constrained permutation as a null model–i.e. a “spin test” [33, 34]. Panels *e*-*j* show analogous plots for cluster centroids 2 and 3.

These results reify, using human imaging data, the observations made using mouse brain networks and suggest that events detected using human fMRI may also be underpinned by modular sub-divisions of the connectome.

## DISCUSSION

Here, we applied edge time series to functional imaging data recorded from anesthetized mice. In agreement with findings made using human imaging data, we find evidence of high-amplitude events. Further, we show that on a per-frame basis events best predict static FC and can be meaningfully grouped into putative co-fluctuation “states”. Lastly, we show that the co-fluctuation patterns expressed during events correspond to highly modular anatomical subgraphs, positing a structural scaffold for events to emerge. This study is one of the first to apply edge-centric network methods to non-human imaging data (mouse). Further, this study adds nuance to and contextualizes analogous observations made using human imaging data while paving the way for future work to investigate origins of high-amplitude BOLD fluctuations using perturbative and possibly invasive experimental techniques.

### Events represent a low-dimensional repertoire of high-amplitude co-fluctuation states that define static FC

Numerous studies have shown that patterns of brain activity and connectivity approximately recur within a given scan session [8, 44–4 These observations are not limited to human fMRI, but have made in other mammals including mice [47–49] and with other imaging modalities, including two-photon and meso-scale calcium imaging [50].

Here, we partition events into non-overlapping clusters using a bespoke hierarchical and recursive variant of modularity maximization. Not only do we find evidence of shared co-fluctuation patterns that recur across time and different mice, we find that they can be meaningfully described at different organizational levels. We focus on the coarsest level, where we detect three large and distinct patterns of high-amplitude co-fluctuation that cross-link well-known anatomical and functional divisions of the mouse brain. Critically, the network organization of static FC is well-explained by these high-amplitude states alone, mirroring findings made using human imaging data [15, 18, 19].

We also examined the activity modes that underlie each of these clusters. Interestingly, they bore a striking resemblance to the results of a recent analysis of high-amplitude mouse activations, in which the authors detected six co-activation patterns (CAPs) that were present in both awake and anesthetized mice [51]. These patterns could be grouped into anti-correlated pairs, such that the spatial patterns of CAPs that correspond to a given pair are approximately anticorrelated with one another.

Notably, the reported CAPs resembled the activity modes that underpinned the event cluster centroids described here (CAPs 3 and 4, 1 and 2n, and 5 and 6 mapped onto clusters 1-3, respectively). This is not a coincidence; mathematically, co-fluctuation matrices are calculated as the product of an activation vector with itself transposed. Consequently, an activation pattern (or the same pattern where the sign of each element is flipped) would generate an identical co-fluctuation matrix [30]. Hence, events can be viewed as a connectivity-based analog of the activation-centric CAPs, and likely explain the parallels between results presented here and in other studies that analyzed the same dataset using CAPs [51, 52].

In the context of these observations and the long history of detecting and tracking network states, our observations suggest that while time-varying functional connectivity is, in principle, a high-dimensional construct, its temporal evolution can be described in terms of relatively few relevant dimensions–i.e. transitions to and from different network states.

We note, however, that the functional/behavioral relevance of these states remains undisclosed. However, given the anesthetized state of the animals, the link to ongoing behavior is tenuous. Rather, they may play a role associated with homeostatic processes [53]. Future studies should investigate this question in greater detail.

### Structural underpinnings of high-amplitude events

Many studies have shown that anatomical connectivity serves as a powerful constraint on both static [1, 2, 54–56], as well as time-varying functional connectivity [57–60]. To date, however, few studies have examined structure-function relationships when function was defined using edge time series (though see [25, 61] for examples).

Here, we study structure-function relationships using an invasively-mapped, directed and weighted connectome through two complementary approaches. First, we show that high-amplitude events result in a division of network nodes into positively and negatively co-fluctuating clusters and that this bipartition is undergirded by highly segregated structural modules. This observation is directly in line with Pope *et al*. [26] and other studies demonstrating that in modular networks, modules easily synchronize [62–64], possibly building to network-wide events. The present study, therefore, represents the first empirical corroboration of Pope *et al*. [26]. Importantly, we verified that this effect was not driven by module size or spatial extent (distance), the parameter combinations needed to detect the bipartition, and was replicated it using human imaging data. Secondly, in a supplementary analysis we tracked moment-to-moment structure-function coupling as the correlation between instantaneous co-fluctuation matrices and structural connectivity. This analysis was similar to Liu *et al*. [61], who predicted co-fluctuation patterns using stylized measures of inter-regional communication capacity [65]. Here, we opted for a simpler metric of coupling based on edge weight correlations, discovering modest coupling across time. Interestingly, however, we found that structure-function coupling was maximized during high-amplitude frames, supporting our previous finding that event co-fluctuations are well-aligned with anatomical connectivity.

Perplexingly, this observation deviates from previous findings. A number of studies using sliding-window methods for tracking time-varying connectivity reported the presence of hyper-connectivity states when global coupling is disproportionately strong [66, 67]. In these studies, hyper-connectivity corresponded to decoupling from the underlying anatomical network [58, 59], with stronger coupling observed in lower-amplitude states. There are, of course, a number of possible explanations. For instance, edge time series and sliding windows measure global amplitude in different ways; whereas sliding window estimates of time-varying connectivity define edge weights as correlation coefficients, effectively placing an upper limit on the global mean connectivity, edge time series have no such bound. Therefore, high-amplitude/hyperconnected time points may not align across methods. This agrees with previous studies that reported only a modest correspondence between time-varying networks estimated using those two techniques [7].

Notably, the observation that modular network structure plays a key role in shaping (near) synchronization patterns is a well documented phenomenon in complex systems and network science [64, 68]. Assortative modules are characterized by dense, recurrent connectivity patterns, allowing for self-excitation of individual modules [69]. On the other hand, the relative segregation of modules from one another ensures that synchronization effects remain localized to subsets of modules, rarely inducing global synchronization stats [62] (though in clustered networks a path to global synchrony becomes possible through the synchronization of the clusters themselves [70]). Though not explicitly tested here, these mechanisms explain observations made in previous simulation studies [25, 26] and align with the empirical findings reported here.

### Events occur in absence of consciousness

Previous analyses of edge time series have largely focused on resting-state or movie-watching conditions (though see [71] for an exception). Although dissimilar, in both cases subjects are conscious throughout the experiment. Here, however, the mice are lightly anesthetized (a common protocol in mouse imaging to reduce in-scanner movement) and unconscious. Nonetheless, each subject’s scan exhibits multiple high-amplitude events.

These findings indicate that consciousness is not a necessary ingredient for events to occur and are therefore in line with Pope *et al*. [26] and Rabuffo *et al*. [25]. Both of these studies show that structurally-constrained dynamics can give rise to neuronal cascades that, when mapped to the infra-slow scale of the fMRI BOLD signal, translate into events.

### Limitations and future directions

This study presents a number of opportunities for future work but also suffers from some limitations. Most notably, to the best of our knowledge, this study represents the first to examine event structure in a model organism. This extension of edge time series is critical; its application to human subjects has left a number of questions unanswered. Namely, it remains unclear why events occur and what are brain/physiological processes that they support. These questions are difficult to answer in the absence of direct, and possibly invasive, measurements and perturbations.

Additionally, future work should consider extending the edge-centric approach from fMRI to other imaging modalities–e.g. widefield calcium imaging [72] or voltage-sensitive indicators [73]. Although the fMRI BOLD signal enjoys a broad agreement with these signals [74], its neuronal provenance is oftentimes unclear and indirect. Relatedly, future studies should also investigate edge time series and events in recordings made in non-mammal brains–e.g. larval zebrafish [75, 76]–where whole-brain activity can be recorded at single-cell resolution [77] and for which anatomical connectivity is mapped at an areal level [78].

There are also a number of limitations associated with this work. For instance, due to the inability to map axonal projections at the level of individual mice (the connectome from Oh *et al*. [40] is a composite of many brains), all structure-function associations were calculated with respect to a single reference connectome. There are many challenges and issues associated with group or consensus-based connectomes in human imaging [43] and, even if such issues are successfully mitigated with invasive tract-tracing, the current analysis makes it impossible to calculate measures of structurefunction correspondence for individual brains.

Yet another potential limitation concerns the null models used to assess the statistical significance of induced modularity. For the mouse data, we compared the observed modularity against an ensemble of equal-sized subgraphs with equal-sized communities but otherwise selected at random. This model does not preserve the geometry of the observed bipartition–i.e. the randomly generated communities are much less spatially compact with no guarantees of spatial contiguity. This is an important feature in brain networks, as both anatomical and functional connection weights are distance dependent [79– Addressing this limitation with mouse data is not straightforward. The accepted approach uses “spin” models to project spatial maps to a spherical surface, rotate the surface randomly, and then project the rotated values back to anatomy [33]. Here, we work with mouse volumetric data, making the implementation of spin tests challenging. To partially address this concern, we replicate our findings using surface-based human imaging data where spin tests easily performed. There, we found that, like the unconstrained permutation test, the observed modularity exceeded that of the null distribution, suggesting that the mouse results may generalize as well. However, future work is needed to confirm that this is the case.

## MATERIALS AND METHODS

### Mouse dataset

#### Mouse resting state fMRI data

All in vivo experiments were conducted in accordance with the Italian law (DL 2006/2014, EU 63/2010, Ministero della Sanitá, Roma) and the recommendations in the Guide for the Care and Use of Laboratory Animals of the National Institutes of Health. Animal research protocols were reviewed and consented by the animal care committee of the Italian Institute of Technology and Italian Ministry of Health. The rsfMRI dataset used in this work consists of n = 19 scans in adult male C57BL/6J mice that are publicly available [85, 86]. Animal preparation, image data acquisition, and image data preprocessing for rsfMRI data have been described in greater detail elsewhere [86]. Briefly, rsfMRI data were acquired on a 7.0-T scanner (Bruker BioSpin, Ettlingen) equipped with BGA-9 gradient set, using a 72-mm birdcage transmit coil, and a four-channel solenoid coil for signal reception. Single-shot BOLD echo planar imaging time series were acquired using an echo planar imaging sequence with the following parameters: repetition time/echo time, 1200/15 ms; flip angle, 30°; matrix, 100 ×100; field of view, 2 × 2 cm2; 18 coronal slices; slice thickness, 0.50 mm; 1500 (n = 19) volumes; and a total rsfMRI acquisition time of 30 min.

Image preprocessing has been previously described in greater detail elsewhere [86]. Briefly, timeseries were despiked, motion corrected, skull stripped and spatially registered to an in-house EPI-based mouse brain template. Denoising and motion correction strategies involved the regression of mean ventricular signal plus 6 motion parameters. The resulting time series were band-pass filtered (0.01-0.1 Hz band) and then spatially smoothed with a Gaussian kernel of 0.5 mm full width at half maximum. After preprocessing, mean regional time-series were extracted for 182 regions of interest (ROIs) derived from a predefined anatomical parcellation of the Allen Brain Institute (ABI, [40, 87]).

#### Mouse Anatomical Connectivity Data

The mouse anatomical connectivity data used in this work were derived from a voxel-scale model of the mouse connectome made available by the Allen Brain Institute [88, 89] (https://data.mendeley.com/datasets/dxtzpvv83k/2). Here, we preserved the directionality of connections–i.e. no symmetrization step was included in the pre-/post-processing pipelines.

Briefly, the mouse structural connectome was obtained from imaging enhanced green fluorescent protein (eGFP)–labeled axonal projections derived 428 viral microinjection experiments, and registered to a common coordinate space [40]. Under the assumption that structural connectivity varies smoothly across major brain divisions, the connectivity at each voxel was modeled as a radial basis kernel-weighted average of the projection patterns of nearby injections [89]. Following the procedure outlined in [88], we re-parcellated the voxel scale model in the same 182 nodes used for the resting state fMRI data, and we adopted normalized connection density (NCD) for defining connectome edges, as this normalization has been shown to be less affected by regional volume than other absolute and/or relative measure of interregional connectivity [90].

### Human imaging dataset

The Human Connectome Project (HCP) 3T dataset [29] consists of structural magnetic resonance imaging (T1w), functional magnetic resonance imaging (fMRI), and diffusion magnetic resonance imaging (dMRI) young adult subjects, some of which are twins. Here we use a subset of the available subjects. These subjects were selected as they comprise the “100 Unrelated Subjects” released by the Connectome Coordination Facility. After excluding data based on completeness and quality control (4 exclusions based on excessive framewise displacement during scanning; 1 exclusion due to software failure), the final subset included 95 subjects (56% female, mean age = 29.29 ± 3.66, age range = 22-36). The study was approved by the Washington University Institutional Review Board and informed consent was obtained from all subjects.

A comprehensive description of the imaging parameters and image preprocessing can be found in [91]. Images were collected on a 3T Siemens Connectome Skyra with a 32-channel head coil. Subjects underwent two T1-weighted structural scans, which were averaged for each subject (TR = 2400 ms, TE = 2.14 ms, flip angle = 8^°^, 0.7 mm isotropic voxel resolution). Subjects underwent four resting state fMRI scans over a two-day span. The fMRI data was acquired with a gradient-echo planar imaging sequence (TR = 720 ms, TE = 33.1 ms, flip angle = 52^°^, 2 mm isotropic voxel resolution, multiband factor = 8). Each resting state run duration was 14:33 min, with eyes open and instructions to fixate on a cross.

Finally, subjects underwent two diffusion MRI scans, which were acquired with a spin-echo planar imaging sequence (TR = 5520 ms, TE = 89.5 ms, flip angle = 78^°^, 1.25 mm isotropic voxel resolution, b-vales = 1000, 2000, 3000 s/mm^2^, 90 diffusion weighed volumes for each shell, 18 b = 0 volumes). These two scans were taken with opposite phase encoding directions and averaged.

Structural, functional, and diffusion images were minimally preprocessed according to the description provided in [91], as implemented and shared by the Connectome Coordination Facility. Briefly, T1w images were aligned to MNI space before undergoing FreeSurfer’s (version 5.3) cortical reconstruction workflow, as part of the HCP Pipeline’s PreFreeSurfer, FreeSurfer, and PostFreeSurfer steps. Functional images were corrected for gradient distortion, susceptibility distortion, and motion, and then aligned to the corresponding T1w with one spline interpolation step. This volume was further corrected for intensity bias and normalized to a mean of 10000. This volume was then projected to the 2mm *32k_fs_LR* mesh, excluding outliers, and aligned to a common space using a multi-modal surface registration [92]. The resultant cifti file for each HCP subject used in this study followed the file naming pattern: `*_Atlas_MSMAll_hp2000_clean.dtseries.nii`. These steps are performed as part of the HCP Pipeline’s fMRIVolume and fMRISurface steps. Each minimally preprocessed fMRI was linearly detrended, band-pass filtered (0.008-0.008 Hz), confound regressed and standardized using Nilearn’s `signal.clean` function, which removes confounds orthogonally to the temporal filters. The confound regression strategy included six motion estimates, mean signal from a white matter, cerebrospinal fluid, and whole brain mask, derivatives of these previous nine regressors, and squares of these 18 terms. Spike regressors were not applied. Following these preprocessing operations, the mean signal was taken at each time frame for each node, as defined by the Schaefer 400 parcellation [93] in *32k_fs_LR* space. Diffusion images were normalized to the mean b0 image, corrected for EPI, eddy current, and gradient non-linearity distortions, and motion, and aligned to subject anatomical space using a boundary-based registration as part of the HCP pipeline’s Diffusion Preprocessing step. In addition to HCP’s minimal preprocessing, diffusion images were corrected for intensity non-uniformity with `N4BiasFieldCorrection` [94]. The Dipy toolbox (version 1.1) [95] was used to fit a multi-shell multi-tissue constrained spherical deconvolution [96] to the data with a spherical harmonics order of 8, using tissue maps estimated with FSL’s `fast` [97]. Tractography was performed using Dipy’s `Local Tracking` module [95]. Multiple instances of probabilistic tractography were run per subject [98], varying the step size and maximum turning angle of the algorithm. Tractography was run at step sizes of 0.25 mm, 0.4 mm, 0.5 mm, 0.6 mm, and 0.75 mm with the maximum turning angle set to 20°. Additionally, tractography was run at maximum turning angles of 10°, 16°, 24°, and 30° with the step size set to 0.5 mm. For each instance of tractography, streamlines were randomly seeded three times within each voxel of a white matter mask, retained if longer than 10 mm and with valid endpoints, following Dipy’s implementation of anatomically constrained tractography [99], and errant streamlines were filtered based on the cluster confidence index [100]. For each tractography instance, streamline count between regions-of-interest were normalized by dividing the count between regions by the geometric average volume of the regions. Since tractography was run nine times per subject, edge values were collapsed across runs. To do this, the weighted mean was taken with weights based on the proportion of total streamlines at that edge. This operation biases edge weights towards larger values, which reflect tractography instances better parameterized to estimate the geometry of each connection.

### Edge time series

Functional connectivity (FC) refers to the magnitude of statistical dependence between activity recorded from distant brain sites. Consider regions *i* and *j* whose activity is denoted by the vectors **x**_*i*_ = [*x*_*i*_(1), …, *x*_*i*_(*T*)] and **x**_*j*_ = [*x*_*j*_(1), …, *x*_*j*_(*T*)]. We can estimate their FC as 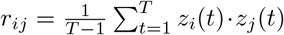, where *z*_*i*_(*t*) and *z*_*j*_(*t*) represent the standardized (z-scored) regional time series. Suppose we omitted the summation in calculating FC. Rather than the correlation coefficient *r*_*ij*_, we would obtain the time series *r*_*ij*_(*t*) = [*z*_*i*_(1) · *z*_*j*_(1), …, *z*_*i*_(*T*) · *z*_*j*_(*t*)]. The elements of this time series have intuitive interpretations; they encode the magnitude, direction, and timing of co-fluctuations between regions *i* and *j*. For instance, *r*_*ij*_(*t*) > 0 if at time *t* regions *i* and *j* both deflect in the same direction with respect to their means–i.e. *sign*(*z*_*i*_(*t*)) = *sign*(*z*_*j*_(*t*)). On the other hand, if *i* and *j* were deflecting in opposite direction, the *r*_*ij*_(*t*) < 0. Relatedly, if at time *t*, the activity of *i* and *j* only slightly deviated from their respective means, the |*r*_*ij*_| ≈ 0. However, if the activity of either region deviates far from its mean, then |*r*_*ij*_| >> 0.

#### Event detection

Co-fluctuation time series have other useful properties. By design, they are an exact decomposition of a functional connection into its time-varying contributions. In previous studies, we found that most frames contribute little to the static connection weight. Rather, FC was driven by a select subset of high-amplitude frames. Across node pairs, these high-amplitude co-fluctuations tended to occur synchronously, giving rise to brain-wide high-amplitude “events”. In previous studies, we detected events by identifying frames where a measure of whole-brain co-fluctuation amplitude was statistically greater than that of a null model. Specifically, we calculated the root mean square (RMS) of all co-fluctuation time series: 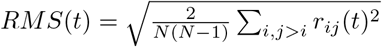. From this time series, we identified its peaks – their amplitude and their timing. We then calculated *RMS* time series using co-fluctuation time series estimated after circularly shifting the regional (parcel) time series. We repeated this procedure 1000 times, building up a null distribution of peak *RMS* values, against which we compared the empirical values using non-parametric statistics. Events were defined as peaks in the intact co-fluctuation time series whose amplitude was statistically greater than that of the null distribution (false discovery rate fixed at 5% and critical *p*-value adjusted accordingly).

### Lin’s concordance

We measured the similarity between co-fluctuation patterns using Lin’s concordance as opposed to the bivariate product-moment correlation. For two patterns with equal means and variances these measures are identical. However, the concordance measure penalizes the similarity if the means differ from one another. For two vectorized co-fluctuation patterns *x* and *y*, concordance is calculated as:

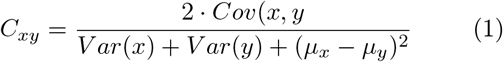

where *Cov*(*x, y*) is the covariance, *V ar*(*x*) is the variance, and *μ*_*x*_ is the mean.

### Modularity heuristic

Many networks exhibit meso-scale or community structure. This implies that they can be meaningfully decomposed into sub-networks referred to a modules or communities. The identity of these sub-networks are usually unknown ahead of time and cannot be determined from visual inspection alone, necessitating algorithmic approaches for estimating nodes’ community assignments. The modularity heuristic is an objective function that maps a network and partition of its nodes into non-overlapping communities to a scalar measure of quality, *Q* [41]. Intuitively, larger values of *Q* are considered “better” partitions.

In more detail, *Q* can be defined as:

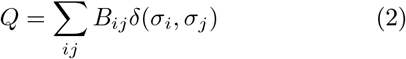

where *B*_*ij*_ is {*i, j*} element of the modularity matrix, *B* = *W* − *P*, where *W* is the observed connectivity matrix and *P* is the connectivity matrix expected under a null model. The function *δ*(*x, y*) is the Kronecker delta and is equal to 1 when *x* = *y* and 0 otherwise. The variable *σ*_*i*_ denotes the community assignment of node *i*. In short, *Q* is calculated as the sum of all within-community elements of the modularity matrix, *B*. It takes on a large value when the observed weights of those connections exceed their expected weights.

The modularity, *Q*, can also be expressed as a sum over communities. Given a partition of nodes into *K* communities, we can write the contribution of community *c* ∈ {1, …, *K*} to the total modularity as *q*_*c*_ = Σ_*i*∈*c,j*∈*c*_ *B*_*ij*_ such that *Q* = Σ_*c*_ *q*_*c*_.

#### Hierarchical and recursive modularity algorithm

Previously we had described an algorithm for recursively applying modularity maximization to obtain a hierarchical partition of a network [18]. The algorithm works as follows. Given a fully-weighted, symmetric, and possibly signed network, we denote its observed connectivity as *C* and define its expected connectivity to be the mean of its upper triangle elements, 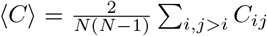. We can then define the modularity matrix, *B* = *C* − ⟨*C*⟩. Using the Louvain algorithm [101], we optimize *Q N*_*iter*_ times and perform consensus clustering on the ensemble of high-quality partitions [102] using a previously described algorithm [103]. This step results in a partition of nodes into *K* communities and the contribution of each community to the total modularity, *q*_*c*_, *c* ∈ {1, …, *K*}.

Each of the *K* communities can be viewed as a “child” of the “parent” network. To obtain a full multi-level and hierarchical description of the network’s communities, we could recursively apply the above procedure to all child networks and subsequently to the children of children and so on, until at the final level every node is its own community. However, this procedure would be computationally expensive, especially for large networks. Moreover, many of the child networks may be poorly defined and not composed of cohesively connected nodes.

Accordingly, we introduce a statistical criterion that prunes branches from the hierarchy. Specifically, after obtaining the consensus partition, we permute consensus community labels to obtain a null distribution for communities’ modularity contributions. Any community whose contribution was consistent with the null distribution was discarded and not sub-divided further, effectively pruning its children from the hierarchical tree. In this manuscript, we used the hierarchical algorithm both to partition static FC into communities at multiple resolutions as well as to cluster high-amplitude events into putative states.

#### Bipartition detection

In a previous study, we showed that co-fluctuation time series induce a bipartition of network nodes into two clusters [30]. One of the clusters corresponds to nodes with positive activations while the other cluster corresponds to those with negative activations. The temporal average of co-assignment matrices obtained from these bipartitions was highly correlated with static FC.

When many co-fluctuation patterns are averaged together, as they are when we estimate event cluster centroids, estimating the bipartition is not straightforward.To obtain such an estimate we resort to data-driven algorithms. Namely, modularity maximization. Specifically, we define a modularity matrix by comparing the observed co-fluctuation matrix against a uniform null model. That is, *B*_*ij*_ = *W*_*ij*_ − *P*, where *P* is a constant and is the same for all {*i, j*}. We optimize the corresponding modularity *N*_*iter*_ = 1000 times and use consensus clustering to obtain a single representation partition. From this partition, we extract the two largest clusters and, by inspection, ensure that they are anticorrelated with one another. We retain this two clusters as an estimate of the bipartition and discard any smaller clusters, assigning nodes in those clusters to a single “non-cluster” label.

#### Induced structural modularity

Given an estimate of bipartition, we wanted to assess whether the two communities were also structurally segregated from one another–i.e. whether the bipartition was modular. To do so, we extracted the subgraph from the structural network–i.e. the connectome–composed of nodes in either of the two clusters. We also extracted the corresponding subgraph from the structural modularity matrix. Here, the modularity matrix was defined as 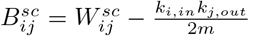, where 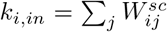 weighted in-degree of node *i* and 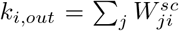 and 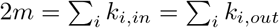.

Suppose that we let *c*^+^ and *c*^*−*^ correspond to the two clusters detected in the bipartition analysis and represent groups of nodes with high-levels of positive and negative activity, respectively. Note that *c*^+^ ∩ *c*^*−*^ = {∅}.Then we can calculate the induced modularity of the bipartition as:

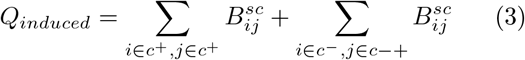

To assess the statistical significance of the induced modularity, we compared the observed modularity against a null distribution generated using permutation tests. For two non-overlapping communities, *c*^+^ and *c*^*−*^, we generated a null distribution by sampling *n*^+^ and *n*^*−*^ at random and recomputing the induced modularity. Note that for the human imaging data, rather than use a random sample, we sampled communities using a spatially constrained “spin test” [33, 34, 104]. To do this, we defined a community vector of length *N*, where *N* is the number of nodes. Elements of *c*^+^ and *c*^*−*^ were assigned values of 1 and 2 respectively while all other elements were equal to 0. The spin test permutes this vector while approximately preserving the spatial contiguity of neural elements. After “spinning” the vector, we extracted the subgraph corresponding to the non-zero elements in the community vector. Thus, this null model assesses whether subgraphs with similar spatial extent, equal size, and equal-sized communities could have generated a similar induced modularity.

## ACKNOWLEDGEMENTS

ER was supported by the Oberlin College Junior Practicum program. RFB acknowledges support from the National Science Foundation (NCS-FO award #2023985). JF acknowledges support from National Institutes of Mental Health (1ZIAMH002783-20).

---

**Figure S1.**
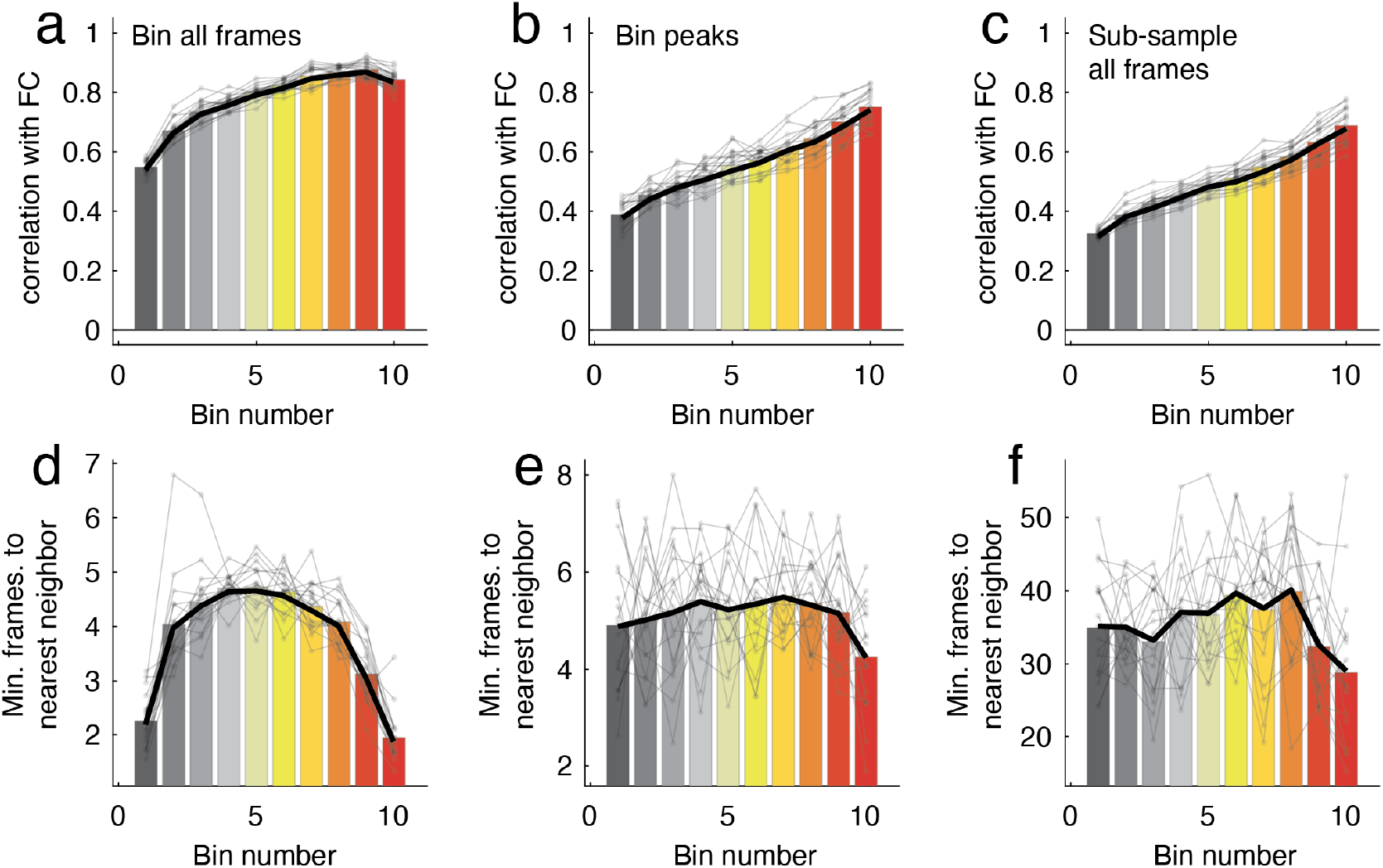
Effect of RMS amplitude and from sampling strategy on predicted FC. In the main text we predicted FC using select subsets of frames. Here, subdivide frames into percentile bins based on their RMS. We explore several binning strategies. (*a*) First we partition all frames from each scan into deciles. As in Cutts *et al*. [16], Ladwig *et al*. [22], we observe a near-monontonic increase in correlation but also find that the second highest amplitdue bin exhibits a marginally greater correspondence with FC. (*b*) The second strategy for binning involves first selecting peak frames of the RMS signal but otherwise divides the frames into RMS deciles. Here, we find a true monotonic increase in RMS across bins. (*c*) We can also use the bins defined in *a* and subsample a number of frames equal to a much smaller number–in this case, the number of peaks per bin from *b*. What explains why *a* exhibits a peak in the second to last bin while strategies *b* and *c* exhibit monotonic increases? Strategy *a* bins all frames. Given the strong autocorrelation in fMRI BOLD data and the fact that events take longer to unfold relative to lower-amplitude peaks [18], the highest bin tends to include many temporally proximal frames. We quantify the relative “nearness” of frames to one another by calculating for each frame assigned to a given bin its nearest neighbor and then averaging those values. When we plot these values for each bin, we find that strategy *a* yields an inverted U-shaped curve, suggesting that the highest (and lowest) amplitude bins are composed of more temporally contiguous frames than middle-amplitude frames and therefore may not effectively sample the entire time series, introducing a bias (see *d*). In contrast, strategies *b* and *c*, which sample peaks and a small number of random samples, exhibit near-uniform nearest neighbor curves, suggesting that if a bias exists, it may be less severe (panels *e* and *f*).

**Figure S2.**
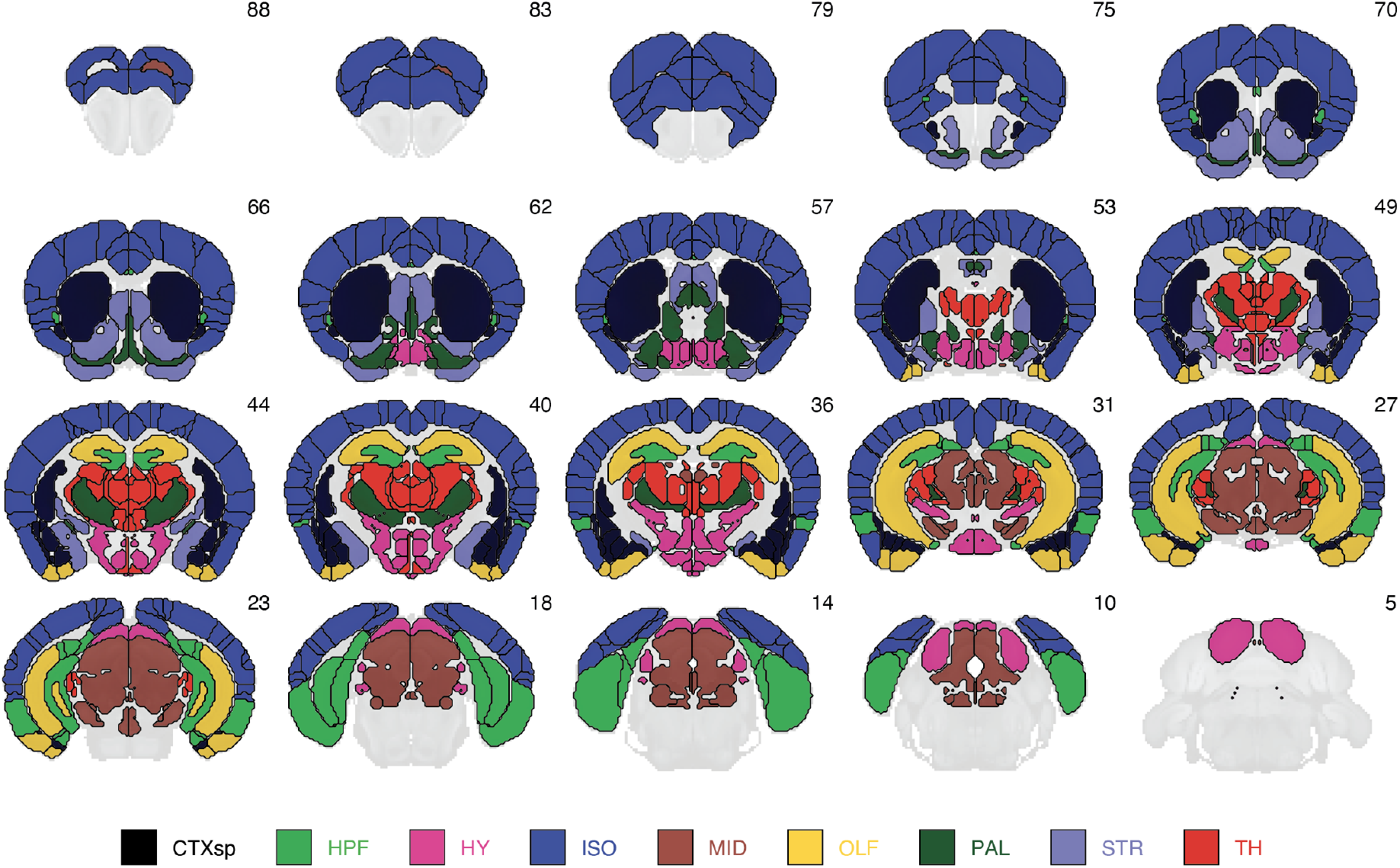
Macroscopic system labels based on anatomy.

**Figure S3.**
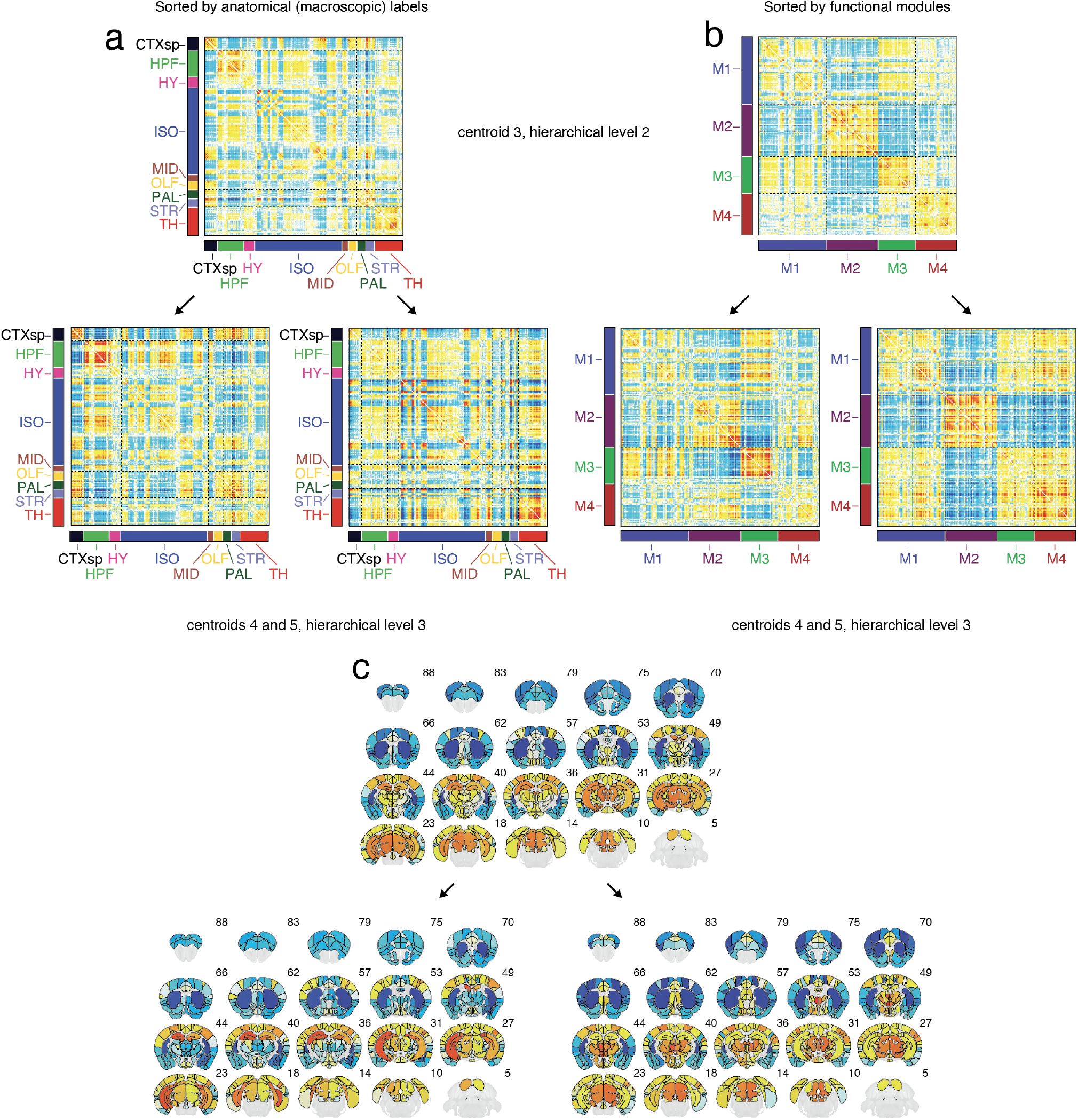
Sub-divisions of event cluster 3. Event cluster 3 in hierarchical level 2 gets subdivided into two clusters at hierarchical level 3 (labeled clusters 4 and 5). (*a*) Cluster centroids ordered by anatomical system labels. (*b*) Cluster centroids ordered by functional systems. (*c*) Leading eigenvectors for each cluster centroid projected onto brain volume.

**Figure S4.**
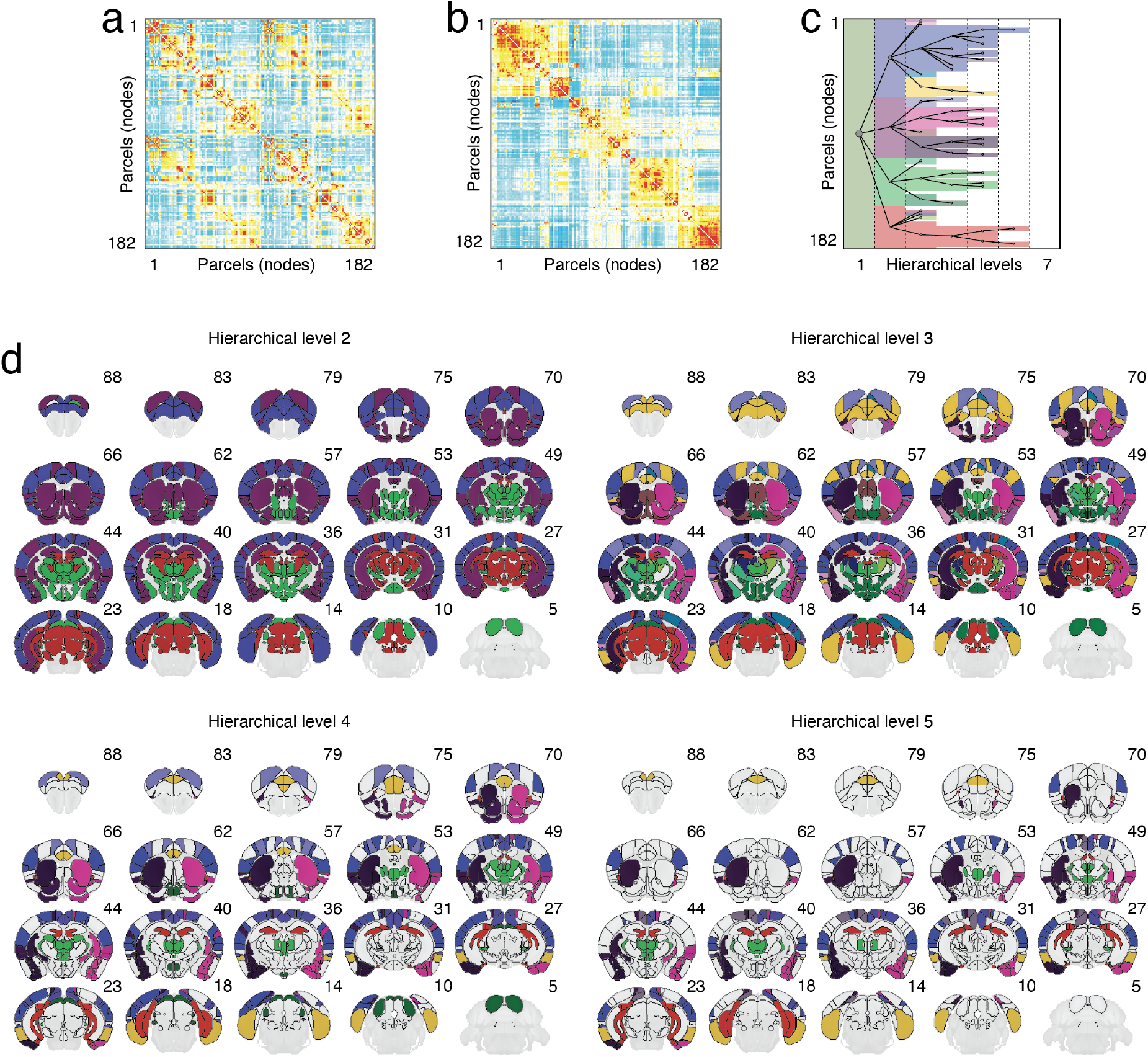
Hierarchical decomposition of static FC into brain systems. (*a*) Unsorted FC matrix. (*b*) Optimally sorted FC matrix. (*c*) Hierarchical community labels and dendrogram. (*d*) Functional system labels at different hierarchical levels.

**Figure S5.**
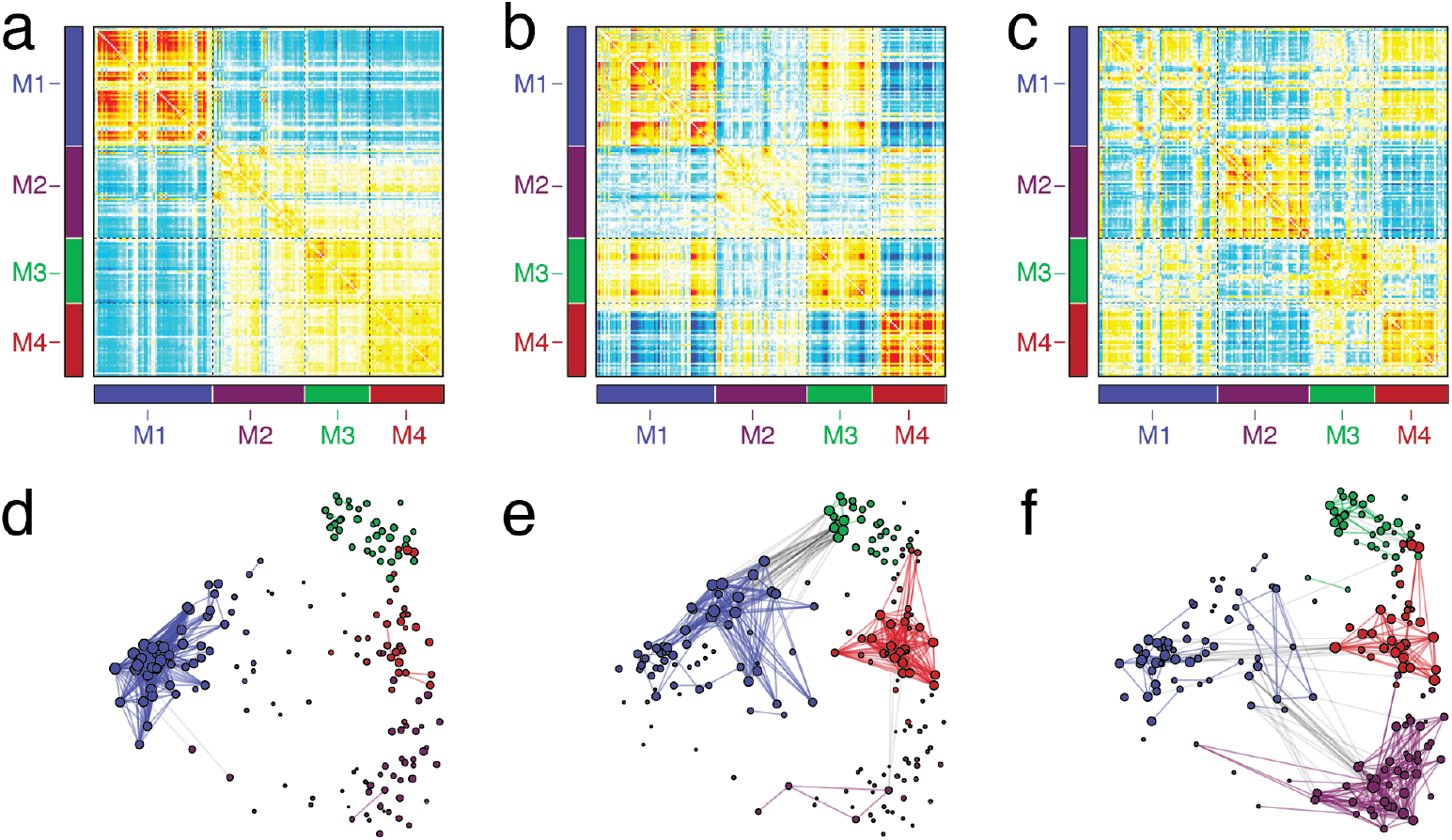
Event co-fluctuation patterns at coarse scale. Panels *a*-*c* depict the same co-fluctuation patterns shown in Fig. 3. However, here we sort rows and columns by functional system labels. Panels *d* -*f* depict co-fluctuation matrices thresholded at 2.5% sparsity. Nodes are colored based on the functional system label. Layout (nodal coordinates) was determined by force-directed algorithm applied to fully-weighted connectome. Note that the aim of this figure is to reinforce the idea that the strongest co-fluctuations (edges) in each cluster centroid tend to fall within functional modules.

**Figure S6.**
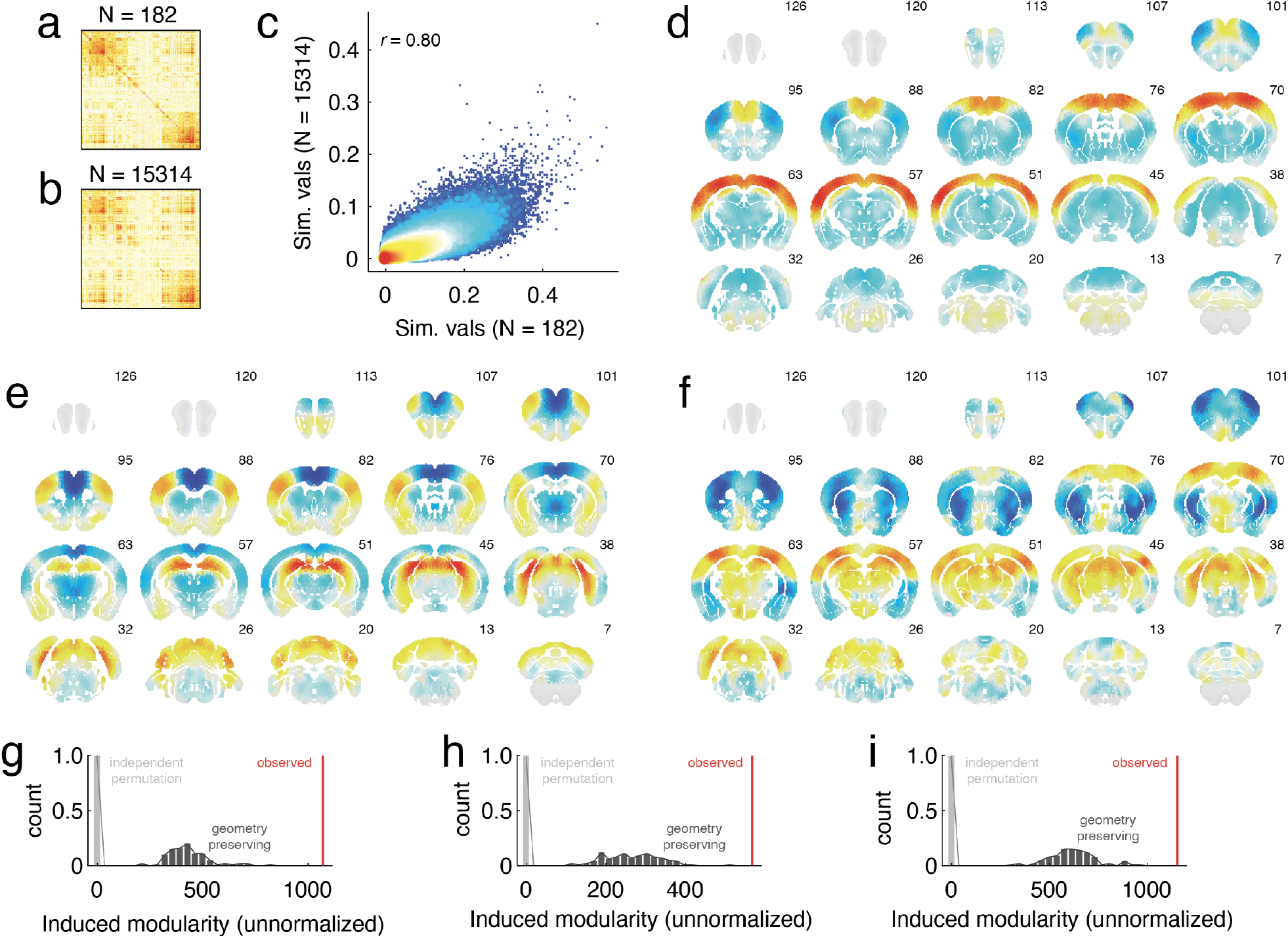
Voxel-level replication of main results. Similarity between pairs of events for parcellated data from main text (*a*) and voxel-wise data (*b*) derived from Coletta *et al*. [88]. (*c*) Scatterplot of upper triangle elements from panels *a* and *b*. Panels *d, e*, and *f* show cluster centroids 1-3 from the main text but at voxel resolution. Panels *d* -*f* show induced modularity of bipartitions compared to permutation-based null model. Note that here, structural connectivity is defined based on the fully-weighted (unthresholded) connectome described by Coletta *et al*. [88].

**Figure S7.**
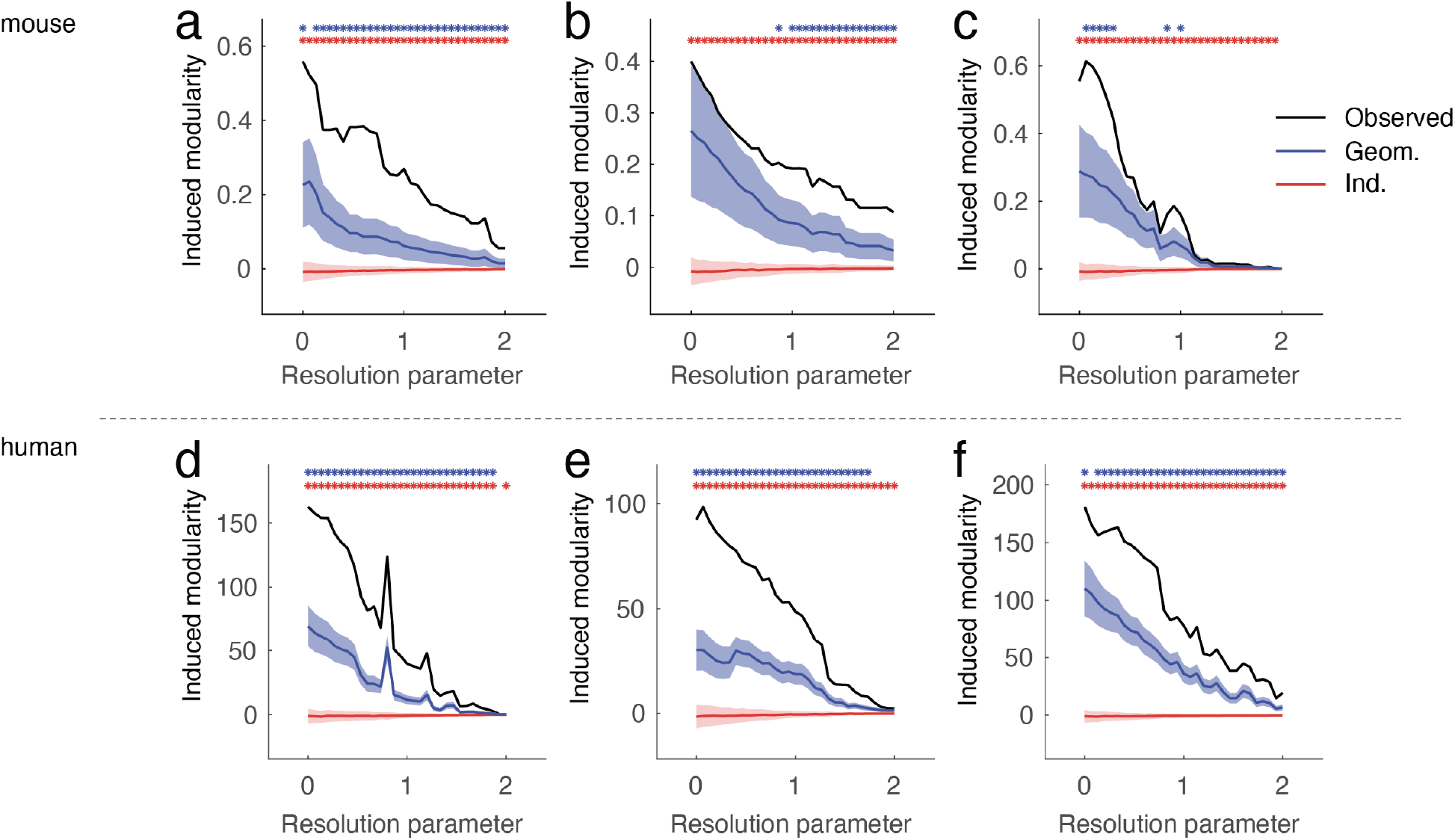
Parametric variation in induced modularity. In the main text we calculated the modularity of a subgraph induced by a bipartition derived from event cluster centroids. Here, we explore the effect of varying the threshold parameter used to define the two clusters. Panels *a*-*c* show the induced modularity of the observed subgraph compared against the modularity estimated under two null models – one that preserves geometry (Geom.) and another that does (Ind.). Blue and red stars indicate statistical significance (critical p-value adjusted to maintain a false discovery rate of 5%). Panels *d* -*f* show analogous plots for the human imaging data.

**Figure S8.**
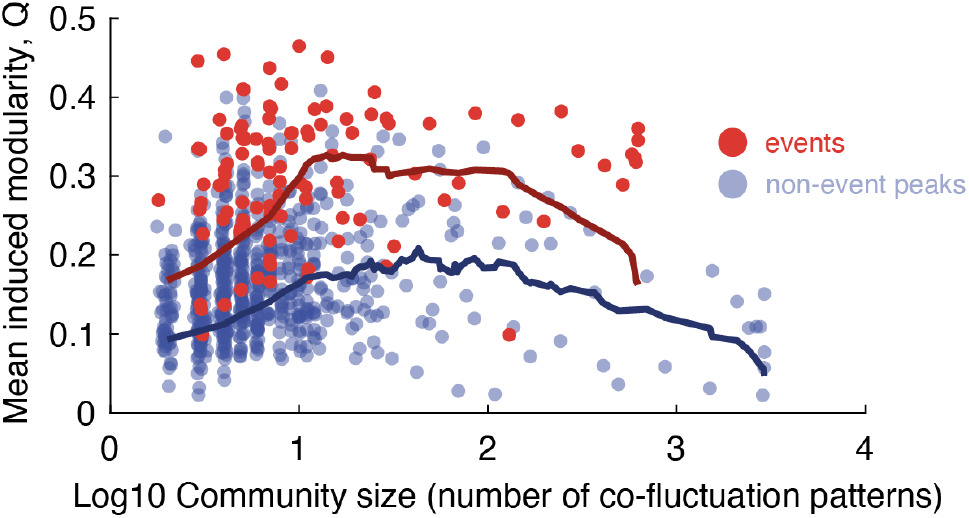
Induced modularity for events *versus* non-event peaks. In the main text we showed that the high-amplitude events are undergirded by a highly modular bipartition of the anatomical network. Here, we show that, the induced modularity of that network is greater than networks of equal size but corresponding to non-event peaks. Specifically, we calculated the mean induced modularity for all subnetworks of size *N*_*s*_. We repeated this procedure for both the events (red) and non-event peaks (blue). We used functional data analysis to compare the two curves. Specifically, we calculated the summed difference between each point. We compared this value against a null model in which we randomly assigned event and non-event centroids to either class. We found that the observed difference in curves was statistically greater than that of the null distribution (*p <* 10^*−*2^).

**Figure S9.**
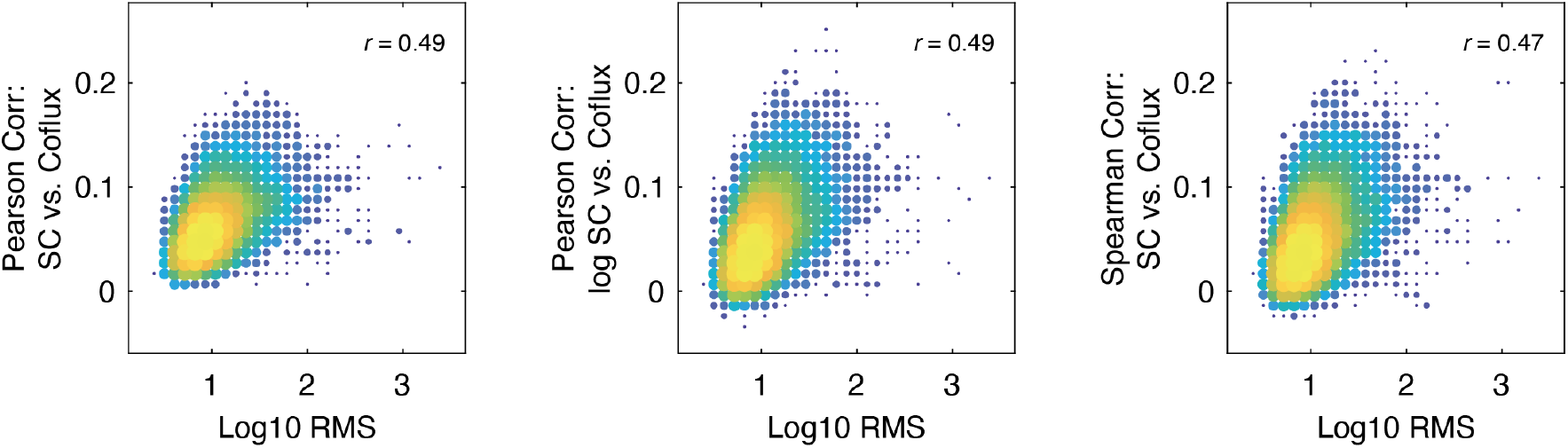
Framewise structure-function correlations scale with RMS. At each frame, we calculated the correlation between SC and the co-fluctuation matrix. We calculated three separate versions of the correlation coefficient. First, we calculated the bivariate product-moment correlation between the raw SC weights and co-fluctuation amplitudes. We also calculated the bivariate correlation using log-transformed SC weights. Finally, we calculated the rank correlation between SC weights and co-fluctuation amplitudes. We compared these frame-resolved structure-function correlation coefficients with the RMS amplitude of global co-fluctuations at each frame. In all cases, we found that structure-function correlation was strongest during high-amplitude frames.

**Figure S10.**
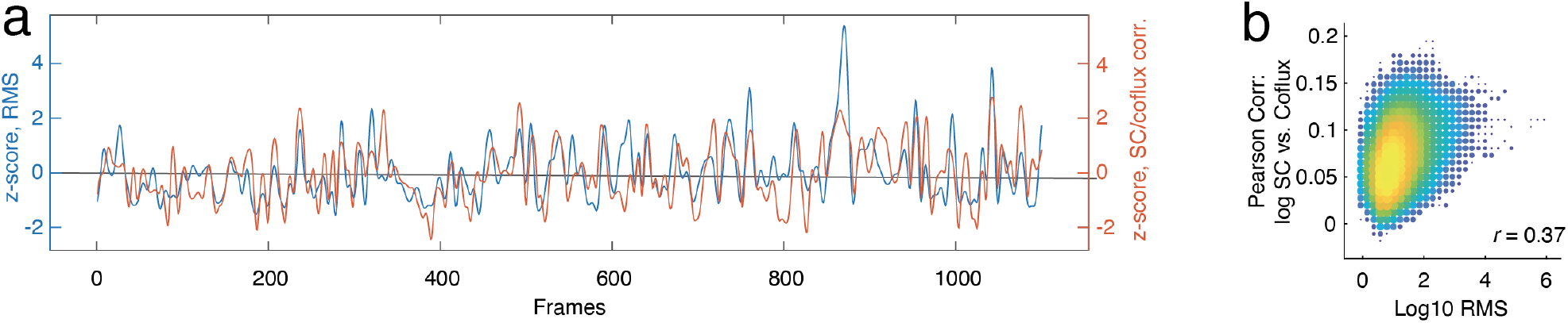
Framewise structure-function correlations scale with RMS using human data from HCP. (*a*) Plot showing example z-scored RMS and correlation coefficient between structural edge weights and corresponding elements of the time-varying co-fluctuation matrix. (*b*) Two-dimensional histogram of the RMS *versus* correlation.

## References

[1] C. J. Honey, R. Kötter, M. Breakspear, and O. Sporns, Proceedings of the National Academy of Sciences 104, 10240 (2007).

[2] C. J. Honey, O. Sporns, L. Cammoun, X. Gigandet, J.-P. Thiran, R. Meuli, and P. Hagmann, Proceedings of the National Academy of Sciences 106, 2035 (2009).

[3] R. M. Hutchison, T. Womelsdorf, E. A. Allen, P. A. Bandettini, V. D. Calhoun, M. Corbetta, S. Della Penna, J. H. Duyn, G. H. Glover, J. Gonzalez-Castillo, et al., Neuroimage 80, 360 (2013).

[4] D. J. Lurie, D. Kessler, D. S. Bassett, R. F. Betzel, M. Breakspear, S. Kheilholz, A. Kucyi, R. Liégeois, M. A. Lindquist, A. R. McIntosh, et al., Network Neuroscience 4, 30 (2020).

[5] R. Hindriks, M. H. Adhikari, Y. Murayama, M. Ganzetti, D. Mantini, N. K. Logothetis, and G. Deco, Neuroimage 127, 242 (2016).

[6] M. Kudela, J. Harezlak, and M. A. Lindquist, NeuroImage 149, 165 (2017).

[7] F. Z. Esfahlani, L. Byrge, J. Tanner, O. Sporns, D. Kennedy, and R. Betzel, bioRxiv (2021).

[8] X. Liu and J. H. Duyn, Proceedings of the National Academy of Sciences 110, 4392 (2013).

[9] X. Liu, C. Chang, and J. H. Duyn, Frontiers in systems neuroscience 7, 101 (2013).

[10] J. M. Shine, O. Koyejo, P. T. Bell, K. J. Gorgolewski, M. Gilat, and R. A. Poldrack, NeuroImage 122, 399 (2015).

[11] F. I. Karahanoğlu and D. Van De Ville, Nature communications 6, 1 (2015).

[12] J. Cabral, D. Vidaurre, P. Marques, R. Magalhães, P. S. Moreira, J. M. Soares, G. Deco, N. Sousa, and M. L. Kringelbach, Scientific reports 7, 1 (2017).

[13] F. Z. Esfahlani, Y. Jo, J. Faskowitz, L. Byrge, D. Kennedy, O. Sporns, and R. Betzel, Proceedings of the National Academy of Sciences (2020).

[14] J. Faskowitz, F. Z. Esfahlani, Y. Jo, O. Sporns, and R. F. Betzel, Nature neuroscience 23, 1644 (2020).

[15] R. F. Betzel, S. A. Cutts, S. Greenwell, J. Faskowitz, and O. Sporns, NeuroImage 252, 118993 (2022).

[16] S. A. Cutts, J. Faskowitz, R. F. Betzel, and O. Sporns, Cerebral Cortex (2022).

[17] L. Sasse, D. I. Larabi, A. Omidvarnia, K. Jung, F. Hoffstaedter, G. Jocham, S. B. Eickhoff, and K. R. Patil, bioRxiv, 2022 (2022).

[18] R. Betzel, S. Cutts, J. Tanner, S. Greenwell, T. Varley, J. Faskowitz, and O. Sporns, bioRxiv (2022).

[19] S. Greenwell, J. Faskowitz, L. Pritschet, T. Santander, E. G. Jacobs, and R. F. Betzel, bioRxiv (2021).

[20] L. Novelli and A. Razi, arXiv preprint arXiv:2106.10631 (2021).

[21] T. Matsui, T. Q. Pham, K. Jimura, and J. Chikazoe, NeuroImage, 118904 (2022).

[22] Z. Ladwig, B. A. Seitzman, A. Dworetsky, Y. Yu, B. Adeyemo, D. M. Smith, S. E. Petersen, and C. Gratton, NeuroImage 260, 119476 (2022).

[23] J. C. Tanner, J. Faskowitz, L. Byrge, D. Kennedy, O. Sporns, and R. Betzel, bioRxiv, 2022 (2022).

[24] G. Levakov, O. Sporns, and G. Avidan, bioRxiv (2022).

[25] G. Rabuffo, J. Fousek, C. Bernard, and V. Jirsa, Eneuro 8(2021).

[26] M. Pope, M. Fukushima, R. Betzel, and O. Sporns, bioRxiv (2021).

[27] J. H. Lee, R. Durand, V. Gradinaru, F. Zhang, Goshen, D.-S. Kim, L. E. Fenno, C. Ramakrishnan, and K. Deisseroth, Nature 465, 788 (2010).

[28] F. Rocchi, C. Canella, S. Noei, D. Gutierrez-Barragan, L. Coletta, A. Galbusera, A. Stuefer, S. Vassanelli, M. Pasqualetti, G. Iurilli, et al., Nature communications 13, 1056 (2022).

[29] D. C. Van Essen, S. M. Smith, D. M. Barch, T. E. Behrens, E. Yacoub, K. Ugurbil, W.-M. H. Consortium, et al., Neuroimage 80, 62 (2013).

[30] O. Sporns, J. Faskowitz, S. Teixera, and R. Betzel, bioRxiv (2020).

[31] J. M. Stafford, B. R. Jarrett, O. Miranda-Dominguez, B. D. Mills, N. Cain, S. Mihalas, G. P. Lahvis, K. M. Lattal, S. H. Mitchell, S. V. David, et al., Proceedings of the National Academy of Sciences 111, 18745 (2014).

[32] J. D. Whitesell, A. Liska, L. Coletta, K. E. Hirokawa, P. Bohn, A. Williford, P. A. Groblewski, N. Graddis, L. Kuan, J. E. Knox, et al., Neuron 109, 545 (2021).

[33] R. D. Markello and B. Misic, NeuroImage, 118052 (2021).

[34] F. Váša and B. Mišić, Nature Reviews Neuroscience 23, 493 (2022).

[35] C. Thomas, Q. Y. Frank, M. O. Irfanoglu, P. Modi, K. S. Saleem, D. A. Leopold, and C. Pierpaoli, Proceedings of the National Academy of Sciences 111, 16574 (2014).

[36] C. Reveley, A. K. Seth, C. Pierpaoli, A. C. Silva, D. Yu, R. C. Saunders, D. A. Leopold, and Q. Y. Frank, Proceedings of the National Academy of Sciences 112, E2820 (2015).

[37] K. H. Maier-Hein, P. F. Neher, J.-C. Houde, M.-A. Côté, E. Garyfallidis, J. Zhong, M. Chamberland, F.-C. Yeh, Y.-C. Lin, Q. Ji, et al., Nature communications 8, 1 (2017).

[38] J.-F. Mangin, P. Fillard, Y. Cointepas, D. Le Bihan, V. Frouin, and C. Poupon, Neuroimage 80, 290 (2013).

[39] M. Reisert, I. Mader, C. Anastasopoulos, M. Weigel, S. Schnell, and V. Kiselev, Neuroimage 54, 955 (2011).

[40] S. W. Oh, J. A. Harris, L. Ng, B. Winslow, N. Cain, S. Mihalas, Q. Wang, C. Lau, L. Kuan, A. M. Henry, et al., Nature 508, 207 (2014).

[41] M. E. Newman and M. Girvan, Physical review E 69, 026113 (2004).

[42] J. B. Burt, M. Helmer, M. Shinn, A. Anticevic, and J. D. Murray, NeuroImage 220, 117038 (2020).

[43] R. F. Betzel, A. Griffa, P. Hagmann, and B. Mišić, Network neuroscience 3, 475 (2019).

[44] E. A. Allen, E. Damaraju, S. M. Plis, E. B. Erhardt, T. Eichele, and V. D. Calhoun, Cerebral cortex 24, 663 (2014).

[45] C. Baldassano, J. Chen, A. Zadbood, J. W. Pillow, U. Hasson, and K. A. Norman, Neuron 95, 709 (2017).

[46] D. Vidaurre, S. M. Smith, and M. W. Woolrich, Proceedings of the National Academy of Sciences 114, 12827 (2017).

[47] J. Grandjean, M. G. Preti, T. A. Bolton, M. Buerge, E. Seifritz, C. R. Pryce, D. Van De Ville, and M. Rudin, Neuroimage 152, 497 (2017).

[48] M. E. Belloy, M. Naeyaert, A. Abbas, D. Shah, V. Vanreusel, J. Van Audekerke, S. D. Keilholz, G. A. Keliris, Van der Linden, and M. Verhoye, Neuroimage 180, 463 (2018).

[49] Y. Ma, C. Hamilton, and N. Zhang, Brain connectivity 7, 1 (2017).

[50] H. Benisty, A. H. Moberly, S. Lohani, D. Barson, R. R. Coifman, G. Mishne, J. A. Cardin, and M. J. Higley, bioRxiv, 2021 (2021).

[51] D. Gutierrez-Barragan, N. A. Singh, F. G. Alvino, L. Coletta, F. Rocchi, E. De Guzman, A. Galbusera, M. Uboldi, S. Panzeri, and A. Gozzi, Current Biology 32, 631 (2022).

[52] D. Fasoli, L. Coletta, D. Gutierrez-Barragan, A. Gozzi, and S. Panzeri, bioRxiv, 2022 (2022).

[53] T. O. Laumann and A. Z. Snyder, Current Opinion in Behavioral Sciences 40, 130 (2021).

[54] Y. Adachi, T. Osada, O. Sporns, T. Watanabe, T. Matsui, K. Miyamoto, and Y. Miyashita, Cerebral cortex 22, 1586 (2012).

[55] R. F. Betzel, J. D. Medaglia, A. E. Kahn, J. Soffer, D. R. Schonhaut, and D. S. Bassett, Nature biomedical engineering 3, 902 (2019).

[56] L. E. Suárez, R. D. Markello, R. F. Betzel, and B. Misic, Trends in Cognitive Sciences (2020).

[57] E. J. Cornblath, A. Ashourvan, J. Z. Kim, R. F. Betzel, R. Ciric, A. Adebimpe, G. L. Baum, X. He, K. Ruparel, T. M. Moore, et al., Communications biology 3, 1 (2020).

[58] R. Liégeois, E. Ziegler, C. Phillips, P. Geurts, F. Gómez, M. A. Bahri, B. T. Yeo, A. Soddu, A. Vanhaudenhuyse, S. Laureys, et al., Brain structure and function 221, 2985 (2016).

[59] M. Fukushima, R. F. Betzel, Y. He, M. P. van den Heuvel, X.-N. Zuo, and O. Sporns, Brain Structure and Function 223, 1091 (2018).

[60] P. Barttfeld, L. Uhrig, J. D. Sitt, M. Sigman, B. Jarraya, and S. Dehaene, Proceedings of the National Academy of Sciences 112, 887 (2015).

[61] Z.-Q. Liu, B. Vazquez-Rodriguez, R. N. Spreng, B. Bernhardt, R. F. Betzel, and B. Misic, bioRxiv (2021).

[62] P. S. Skardal and J. G. Restrepo, Physical Review E 85, 016208 (2012).

[63] P. N. McGraw and M. Menzinger, Physical Review E 72, 015101 (2005).

[64] L. M. Pecora, F. Sorrentino, A. M. Hagerstrom, T. E. Murphy, and R. Roy, Nature communications 5, 4079 (2014).

[65] A. Avena-Koenigsberger, B. Misic, and O. Sporns, Nature Reviews Neuroscience 19, 17 (2018).

[66] A. Zalesky, A. Fornito, L. Cocchi, L. L. Gollo, and M. Breakspear, Proceedings of the National Academy of Sciences 111, 10341 (2014).

[67] R. M. Hutchison, T. Womelsdorf, J. S. Gati, S. Everling, and R. S. Menon, Human brain mapping 34, 2154 (2013).

[68] A. Arenas and A. Diaz-Guilera, The European Physical Journal Special Topics 143, 19 (2007).

[69] Y. S. Cho, T. Nishikawa, and A. E. Motter, Physical review letters 119, 084101 (2017).

[70] F. Sorrentino, L. M. Pecora, A. M. Hagerstrom, T. E. Murphy, and R. Roy, Science advances 2, e1501737 (2016).

[71] J. Rasero, R. Betzel, A. I. Sentis, T. E. Kraynak, P. J. Gianaros, and T. Verstynen, bioRxiv, 2021 (2021).

[72] E. M. Lake, X. Ge, X. Shen, P. Herman, F. Hyder, J. A. Cardin, M. J. Higley, D. Scheinost, X. Papademetris, M. C. Crair, et al., Nature Methods 17, 1262 (2020).

[73] M. H. Mohajerani, A. W. Chan, M. Mohsenvand, J. LeDue, R. Liu, D. A. McVea, J. D. Boyd, Y. T. Wang, M. Reimers, and T. H. Murphy, Nature neuroscience 16, 1426 (2013).

[74] H. Vafaii, F. Mandino, G. Desrosiers-Gregoire, D. O’Connor, X. Shen, X. Ge, P. Herman, F. Hyder, X. Papademetris, M. Chakravarty, et al., bioRxiv, 2022 (2022).

[75] R. F. Betzel, Network Neuroscience 4, 234 (2020).

[76] A. S. Blevins, D. S. Bassett, E. K. Scott, and G. C. Vanwalleghem, Network Neuroscience 6, 1125 (2022).

[77] X. Chen, Y. Mu, Y. Hu, A. T. Kuan, M. Nikitchenko, O. Randlett, A. B. Chen, J. P. Gavornik, H. Sompolinsky, F. Engert, et al., Neuron 100, 876 (2018).

[78] M. Kunst, E. Laurell, N. Mokayes, A. Kramer, F. Kubo, A. M. Fernandes, D. Förster, M. Dal Maschio, and H. Baier, Neuron 103, 21 (2019).

[79] R. F. Betzel and D. S. Bassett, Proceedings of the National Academy of Sciences 115, E4880 (2018).

[80] F. Z. Esfahlani, M. A. Bertolero, D. S. Bassett, and R. F. Betzel, NeuroImage 211, 116612 (2020).

[81] J. Stiso and D. S. Bassett, Trends in cognitive sciences 22, 1127 (2018).

[82] M. Ercsey-Ravasz, N. T. Markov, C. Lamy, D. C. Van Essen, K. Knoblauch, Z. Toroczkai, and H. Kennedy, Neuron 80, 184 (2013).

[83] P. E. Vértes, A. F. Alexander-Bloch, N. Gogtay, J. N. Giedd, J. L. Rapoport, and E. T. Bullmore, Proceedings of the National Academy of Sciences 109, 5868 (2012).

[84] S. Oldham, B. D. Fulcher, K. Aquino, A. Arnatke-vičiūtė, C. Paquola, R. Shishegar, and A. Fornito, Science advances 8, eabm6127 (2022).

[85] J. Grandjean, C. Canella, C. Anckaerts, G. Ayranci, S. Bougacha, T. Bienert, D. Buehlmann, L. Coletta, D. Gallino, N. Gass, et al., Neuroimage 205, 116278 (2020).

[86] D. Gutierrez-Barragan, M. A. Basson, S. Panzeri, and A. Gozzi, Current Biology 29, 2295 (2019).

[87] Q. Wang, S.-L. Ding, Y. Li, J. Royall, D. Feng, P. Lesnar, N. Graddis, M. Naeemi, B. Facer, A. Ho, et al., Cell 181, 936 (2020).

[88] L. Coletta, M. Pagani, J. D. Whitesell, J. A. Harris, B. Bernhardt, and A. Gozzi, Science Advances 6, eabb7187 (2020).

[89] J. E. Knox, K. D. Harris, N. Graddis, J. D. Whitesell, H. Zeng, J. A. Harris, E. Shea-Brown, and S. Mihalas, Network Neuroscience 3, 217 (2018).

[90] M. Rubinov, R. J. Ypma, C. Watson, and E. T. Bullmore, Proceedings of the National Academy of Sciences 112, 10032 (2015).

[91] M. F. Glasser, S. N. Sotiropoulos, J. A. Wilson, T. S. Coalson, B. Fischl, J. L. Andersson, J. Xu, S. Jbabdi, M. Webster, J. R. Polimeni, et al., Neuroimage 80, 105 (2013).

[92] E. C. Robinson, S. Jbabdi, M. F. Glasser, J. Andersson, G. C. Burgess, M. P. Harms, S. M. Smith, D. C. Van Essen, and M. Jenkinson, Neuroimage 100, 414 (2014).

[93] A. Schaefer, R. Kong, E. M. Gordon, T. O. Laumann, X.-N. Zuo, A. J. Holmes, S. B. Eickhoff, and B. T. Yeo, Cerebral cortex 28, 3095 (2018).

[94] N. J. Tustison, B. B. Avants, P. A. Cook, Y. Zheng, A. Egan, P. A. Yushkevich, and J. C. Gee, IEEE transactions on medical imaging 29, 1310 (2010).

[95] E. Garyfallidis, M. Brett, B. Amirbekian, A. Rokem, S. Van Der Walt, M. Descoteaux, and I. Nimmo-Smith, Frontiers in neuroinformatics 8, 8 (2014).

[96] J.-D. Tournier, F. Calamante, and A. Connelly, Neuroimage 35, 1459 (2007).

[97] Y. Zhang, M. Brady, and S. Smith, IEEE transactions on medical imaging 20, 45 (2001).

[98] H. Takemura, C. F. Caiafa, B. A. Wandell, and F. Pestilli, PLoS computational biology 12, e1004692 (2016).

[99] R. E. Smith, J.-D. Tournier, F. Calamante, and A. Connelly, Neuroimage 62, 1924 (2012).

[100] K. M. Jordan, B. Amirbekian, A. Keshavan, and R. G. Henry, Journal of Neuroimaging 28, 64 (2018).

[101] V. D. Blondel, J.-L. Guillaume, R. Lambiotte, and E. Lefebvre, Journal of statistical mechanics: theory and experiment 2008, P10008 (2008).

[102] A. Lancichinetti and S. Fortunato, Scientific reports 2, 1 (2012).

[103] R. F. Betzel, M. A. Bertolero, E. M. Gordon, C. Gratton, N. U. Dosenbach, and D. S. Bassett, Neuroimage 202, 115990 (2019).

[104] F. Váša, J. Seidlitz, R. Romero-Garcia, K. J. Whitaker, G. Rosenthal, P. E. Vértes, M. Shinn, A. Alexander-Bloch, P. Fonagy, R. J. Dolan, et al., Cerebral Cortex 28, 281 (2018).

